# The essential transcriptional regulator HDP1 drives expansion of the inner membrane complex during early sexual differentiation of malaria parasites

**DOI:** 10.1101/2020.10.26.352583

**Authors:** Riward Campelo Morillo, Xinran Tong, Wei Xie, Steven Abel, Lindsey M. Orchard, Wassim Daher, Dinshaw J. Patel, Manuel Llinás, Karine G. Le Roch, Björn F.C. Kafsack

## Abstract

Transmission of Plasmodium *falciparum* and other malaria parasites requires their differentiation from asexual blood stages into gametocytes, the non-replicative sexual stage necessary for transmission to the mosquito vector. This transition involves changes in gene expression and chromatin reorganization that result in the activation and silencing of stage-specific genes. However, the genomes of malaria parasites have been noted for their dearth of transcriptional and chromatin regulators and the molecular mediators of these changes remain largely unknown. We recently identified <u>H</u>omeo<u>D</u>omain <u>P</u>rotein 1 (HDP1) as a DNA-binding protein that enhances the expression of key genes that are critical for early sexual differentiation. The discovery of a homeodomain-like DNA-binding protein marks a new class of transcriptional regulator in malaria parasites outside of the better-characterized ApiAP2 family. In this study, we show that HDP1 facilitates the necessary upregulation of inner membrane complex components during early gametocytogenesis that gives *P. falciparum* gametocytes they characteristic shape and is required for gametocyte maturation and parasite transmission.

## INTRODUCTION

To complete its life cycle, *Plasmodium falciparum*, the most widespread and virulent of the human malaria parasites, must repeatedly differentiate into unique cell types that are able to access and exploit specialized niches within their human and mosquito hosts. One of these key developmental transitions occurs during the parasite’s blood stage. Asexual blood-stages maintain a persistent infection through continuous lytic replication within erythrocytes but are not infectious to the mosquito vector, and therefore cannot mediate transmission to the next human host. Infection of the vector requires asexual blood stages to differentiate into non-replicating, male and female gametocytes that can infect the mosquito once taken up during a blood meal.

All differentiation requires the repression and activation of genes that underlie the specific phenotypes of the origin and destination cell types, respectively. To ensure complete commitment to one cell type or another, these transitions often involve a bistable switch that controls the activity of a single master regulator at the top of the transcriptional cascade that underlies the differentiation program (*1-4*). Upon activation, the master regulator initiates the broader downstream changes in gene expression by altering the expression of additional transcriptional regulators and changing their access to cell type-specific promoters via chromatin re-organization.

Recent work has found that this paradigm also applies in malaria parasites, where the initiation of sexual differentiation is controlled by bistable expression of a master regulator, the transcription factor AP2-G (*5, 6*). During asexual replication the *ap2-g* locus is silenced by heterochromatin (*5, 7, 8*) but, due to the presence of AP2-G binding sites within its own promoter region, incomplete repression of *ap2-g* in individual cells can trigger a transcriptional feedback that drives its expression to high levels, thereby locking cells into the sexual differentiation gene expression program (*5, 9, 10*). Under conditions that impair heterochromatin maintenance, this feedback loop is activated in a larger fraction of cells, thus increasing the frequency of sexual differentiation (*8, 11, 12*).

Commitment to this sexual differentiation program involves substantial changes in genes expression and re-distribution of heterochromatin during the early stages of gametocyte development (*13, 14*). While AP2-G is critical for the initiation of sexual differentiation, it is expressed only during a small window that begins with sexually committed schizonts and ends after the first 48 hours of gametocyte development (*10, 15*). This means that many of the expression changes during the prolonged process of gametocyte development depend on additional transcriptional regulators and is consistent with our previous observations that AP2-G upregulates a number of putative transcription factors and chromatin remodeling enzymes (*9, 10*).

While most species of malaria parasites form spherical gametocytes that mature in 2-6 days, sexual differentiation in *P. falciparum* takes 12-14 days, and produces gametocytes with the characteristic falciform morphology that give the parasite its name. Unsurprisingly, this prolonged maturation is accompanied by a wide array of gene expression changes (*16-19*). However, relatively little is known about the transcriptional regulators downstream of AP2-G that mediate these changes. Compared to other single-celled eukaryotes, DNA-binding proteins are notably underrepresented in the genomes of malaria parasites (*20*). Most belong to the ApiAP2 family (*20*), but only a small number have been shown to function specifically during gametocyte development of *P. falciparum* (*21*). In the rodent malaria parasite *P. berghei*, PbAP2-G2 functions as a transcriptional repressor of asexual-specific gene expression (*6, 22, 23*), while PbAP2-FG (PyAP2-G3 in *P. yoelii*) was shown to mediate upregulation of female-specific transcripts (*24, 25*).

In this study, we identify HDP1, a previously uncharacterized DNA-binding protein that is absent from asexual blood stages and first expressed during early sexual differentiation. The development of HDP1-deficient gametocytes arrests at the Stage I to Stage II transition and ends in loss of viability. Analysis of gene expression and HDP1-binding shows that this protein functions as a positive transcriptional regulator of genes essential for gametocyte development, including genes that are critical for the expansion of the inner membrane complex (IMC) that gives *P. falciparum* gametocytes their characteristic shape.

## RESULTS

### *hdp1* is essential for gametocyte development

Searching for possible regulators of gene expression during *P. falciparum* sexual differentiation, we identified <u>H</u>omeo<u>D</u>omain-like <u>P</u>rotein 1 (*hdp1*, PF3D7_1466200) encoding a 3078 amino acid protein with a C-terminal homeodomain-like domain, containing a helix-turn-helix structural motif commonly involved in DNA binding (fig. S1A) (*26*). Syntenic orthologs of *hdp1* could be readily identified in other malaria parasites with homology to the homeodomain-like domain also found among the coccidia but apparently absent from other apicomplexan clades (fig. S1B). Analysis of *hdp1* transcript levels in *P. falciparum* blood stages by qRT-PCR detected only minimal expression in cultures of asexual blood stages, which always contain small number of gametocytes, but showed substantial upregulation during the early stages of gametocytogenesis (fig. S1C). AP2-G, the transcriptional master switch that controls the initiation of the sexual differentiation gene expression program (*5, 9, 10, 27*), binds at two sites located upstream of *hdp1* in early gametocytes (*10*), consistent with our hypothesis that AP2-G activates additional regulators of gene expression during early gametocytogenesis.

To determine HDP1’s subcellular localization, we inserted an N-terminal HaloTag at the endogenous *hdp1* locus (fig. S2A) to avoid possible interference with the predicted DNA-binding domain (DBD) located at the very C-terminus. As expected, based on transcript abundance data, no Halo-tagged protein was detected in asexual stages. However, when we attempted to determine Halo-HDP1 levels in the sexual stages, we found that *halo-hdp1* cultures were unable to produce the characteristic crescent shapes of maturing *P. falciparum* gametocytes (Fig. 1A-B). Subsequent tagging at the HDP1 C-terminus with either GFP or a triple Ty1 epitope tag (*hdp1-gfp* and *hdp1-Ty1*, fig. S2B-C), yielded parasite lines that produce gametocytes indistinguishable in numbers and morphology from the wildtype parent (Fig. 1A-B), despite the proximity to the putative DNA-binding domain. To test whether N-terminal tagging resulted in a loss of HDP1 function, we generated a Δ*hdp1* knockout line for comparison by replacing 1.4 kb at the 5’ end of the *hdp1* locus with a selectable marker cassette (fig. S2D). The resulting *Δhdp1* parasites exhibited no discernible change in phenotype in asexual blood-stages, but like the *halo-hdp1* parasites, were unable to form viable mature gametocytes (Fig. 1D-F). More detailed analyses using synchronous induction of gametocytogenesis found that both *halo-hdp1* and Δ*hdp1* have sexual commitment rates comparable to the NF54 parent (fig. S3) but are unable to complete gametocyte development (Fig. 1A-C).

**Fig. 1:**
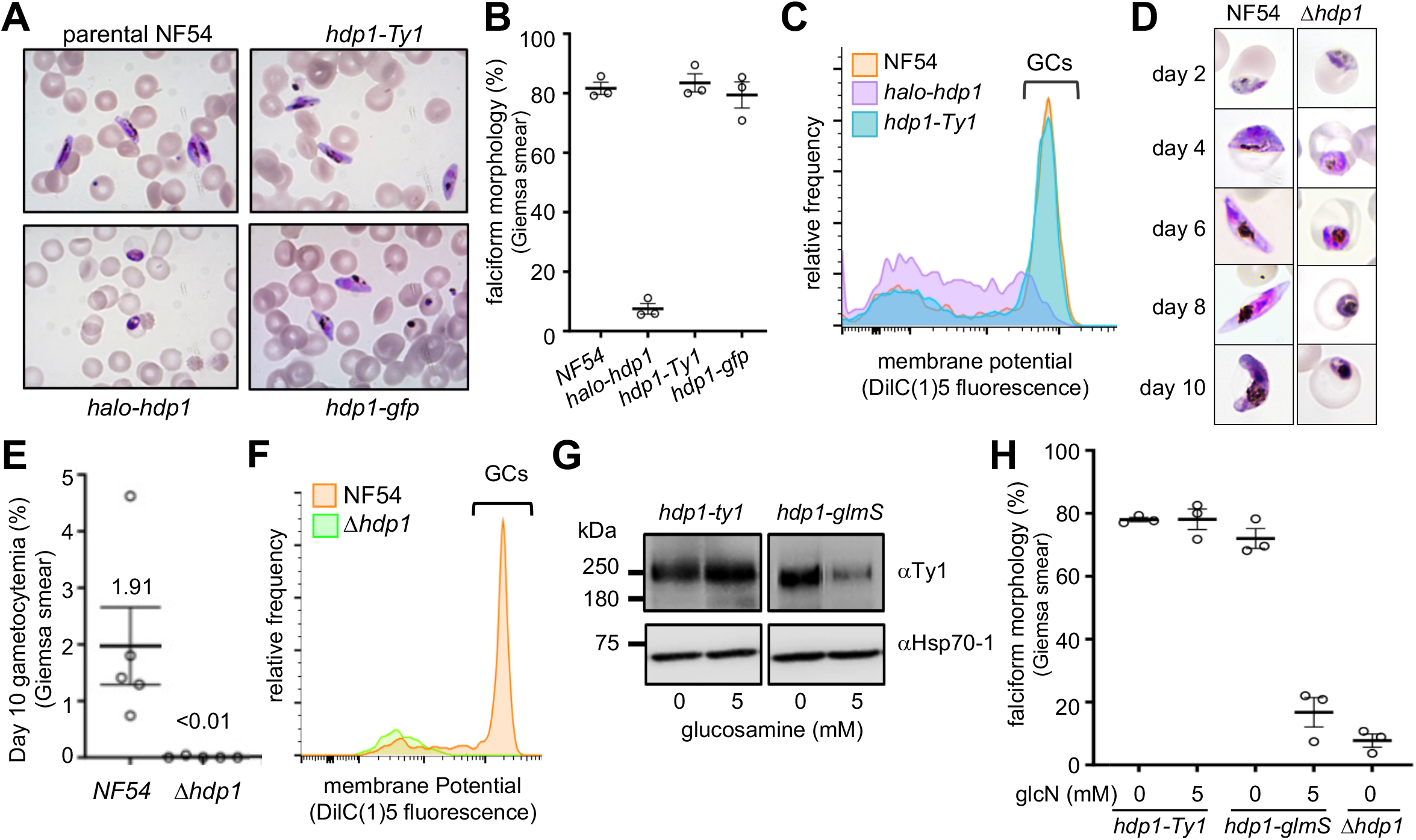
Loss of HDP1 function disrupts gametocyte maturation. **(A-C)** N-terminal tagging of the endogenously encoded HDP1 (*halo-hdp1*) blocked maturation of gametocytes (GCs) while C-terminal tagging of the endogenous locus with either GFP (*hdp1-gfp*) or a triple Ty1 (*hdp1-Ty1*) epitope had no effect on gametocyte morphology or viability, as determined by membrane potential staining with DilC(1)-5 on day 5 of gametocyte maturation. **(D-F)** Targeted disruption of the *hdp1* locus (*Δhdp1*) blocked formation of late gametocytes. Graphs show mean ± s.e.m of n=3-5 **(G)** Glucosamine-inducible knockdown of HDP1 in *hdp1-Ty1* and *hdp1-glmS* day 5 gametocytes. Representative of n=3. **(H)** Percentage of falciform day 5 gametocytes in response to 5 mM glucosamine (GlcN). All Images and flow cytometry plots are representative of n=3.

As the length of the *hdp1* coding sequence made genetic complementation infeasible, we generated inducible HDP1 knockdown parasites by inserting a triple Ty1 epitope tag followed by the autocatalytic *glmS* ribozyme at the 3’ end of the endogenous *hdp1* coding sequence (*hdp1-glmS*, fig. S2E) (*28*). In the absence of glucosamine, the resulting *hdp1-glmS* gametocytes expressed HDP1 protein at levels comparable to *hdp1-Ty1* parasites lacking the ribozyme and produced gametocytes that were indistinguishable from wild-type in both number and morphology (fig. S4). Supplementation of the culture medium with 5 mM glucosamine during the first 5 days of gametocyte development had no discernible effect on *hdp1-Ty1* parasites but diminished HDP1 expression by 70% and reduced the number of falciform gametocytes by 80% in *hdp1-glmS* parasites, recapitulating the phenotype of the *halo-hdp1* and Δ*hdp1* lines (Fig. 1G-H). Since *hdp1* transcript levels remain relatively constant during gametocyte development (fig. S1C), we wanted to test whether *hdp1* transcripts were required throughout gametocyte maturation. Knockdown of *hdp1* levels with glucosamine reduced gametocyte maturation prior to day 5 but not thereafter, indicating that sufficient HDP1 protein had been produced by that time to support gametocyte maturation (fig. S4).

### HDP1 is a chromatin-associated nuclear protein expressed in gametocytes

An earlier study had reported a variety of subcellular localizations based on antibodies raised against a low-complexity region of HDP1 (see fig. S1A), including export to the erythrocyte membrane of early gametocytes (*29*). This was surprising given the absence of a signal peptide, the presence of two predicted nuclear localization signals (fig. S1A), and localization of the *Toxoplasma gondii* ortholog to the nucleus of tachyzoites (fig. S5). To resolve this apparent disagreement, we carried out live-cell and immunofluorescence microscopy of *hdp1-gfp* and *hdp1-Ty1* gametocytes, respectively. In both lines, HDP1 localized exclusively to the gametocyte nucleus (Fig. 2A-B), and we were unable to replicate the localization(s) described in the earlier study. According to its corresponding author, the antisera used in that study are unfortunately no longer available, precluding a direct comparison. HDP1 protein was undetectable in asexual blood stages by both microscopy and western blotting (fig. S6, Fig. 2C) but showed increasing expression from day 2 of gametocytogenesis (Stage I-II) onward, reaching maximal levels by day 5 (Stage III) that remained steady until day 8 (Stage IV) (Fig. 2C). Analysis of subcellular compartments from day 5 *hdp1-Ty1* gametocytes found HDP1 almost exclusively in the nuclear fraction, with about 70% resistant to solubilization up to 600 mM NaCl, indicating a tight association with chromatin (Fig. 2D) and validating HDP1 as a nuclear protein. We were unable to detect HDP1 expression above background in asexual blood stages by either Western blot or immunofluorescence microscopy (Fig. 2C, fig. S6)

**Fig. 2:**
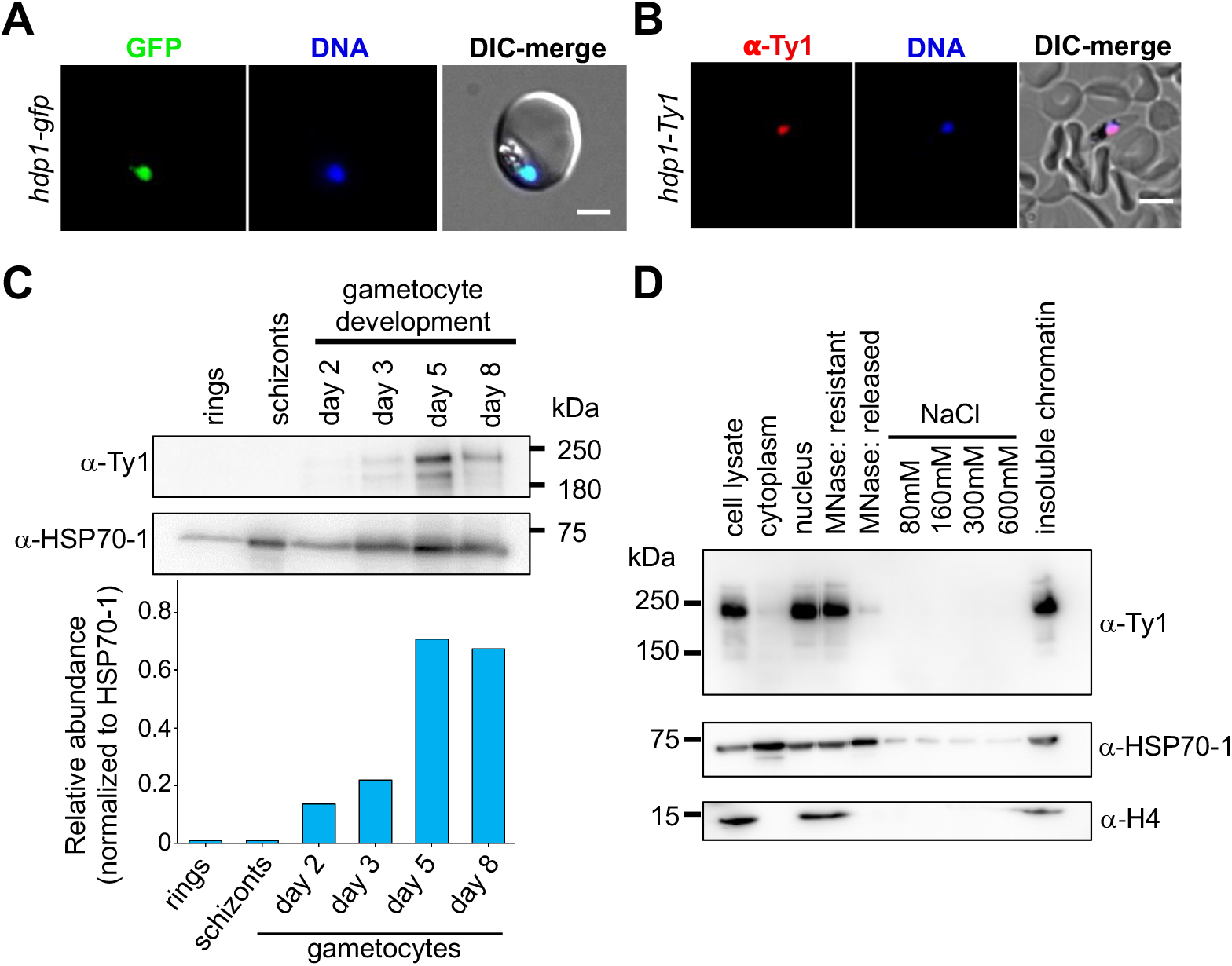
HDP1 is a chromatin-associated protein nuclear protein expressed in gametocytes. **(A)**Live-cell fluorescence microscopy of *hdp1-gfp* gametocytes on day 5 of maturation stained with the DNA dye Hoechst33342 (blue). Scale Bar: 3 µm. Representative of n=2. **(B)** Immunofluorescence microscopy of *hdp1-Ty1* gametocytes on day 5 of maturation co-stained with anti-Ty1 antibodies (red) and Hoechst33342 (blue). Scale Bar: 5 µm. Representative of n=2. **(C)** Western blotting of parasite lysates in asexual stages and during gametocyte maturation shows HDP1 is expressed during the stages of gametocytogenesis. Representative of n=3. **(D)** Western blot of cytoplasmic and nuclear extracts of hdp1-Ty1 gametocytes on day 5 of maturation stained with antibodies against the Ty1 epitope tag, the histone H4, and HSP70-1. Representative of n=3

### Loss of HDP1 leads to dysregulation of gene expression in early gametocytes

Based on HDP1’s nuclear localization and chromatin binding capacity, we wanted to test whether it plays a role in the regulation of gene expression during early gametocytogenesis. In order to identify changes in expression that may be responsible for the aberrant development of Δ*hdp1* gametocytes rather than looking at the consequences of the subsequent loss of viability, we decided to analyze the transcriptome of Δ*hdp1* and parental NF54 on Day 2 of gametocytogenesis, when HDP1 is first detectable in wild-type gametocytes but before any change in viability or morphology occurs in Δ*hdp1* gametocytes. (Fig. 1D, fig. S3B, Data sets 1-2). Global comparison of transcript abundances found that most genes were expressed at similar levels, including canonical markers of early gametocytes such as *pfs16* and *gexp5* (Fig. 3A). This confirms our earlier observation that HDP1-deficient parasites initiate gametocyte development at wild-type rates and that changes in gene expression are not due to change in viability at this point. However, when compared to its parent line, Δ*hdp1* gametocytes had significant reductions in the transcript levels of 156 genes and increased levels of 103 genes. Reassuringly, *hdp1* showed the greatest decrease in expression. Gene set enrichment analysis (GSEA) found that transcripts encoding components of the IMC were significantly over-represented among the down-regulated genes (Fig. 3B-C, Data set 3) while transcripts from heterochromatin-associated multi-copy gene families were significantly over-represented among up-regulated genes (Fig. 3D), including members of the *var, rifin, stevor*, and *PHISTa/b/c* gene families (Fig. 3E, fig. S7A-C). Several of these heterochromatin-silenced families exhibit transcriptional variation due to expression switching between members but this does not lead to the broad upregulation across all family members we observed here. Due to their general lack of expression in wild-type cells, the observed fold-changes for these genes were often substantial but the absolute increase was generally small yet nevertheless significantly above their levels in wild-type cells. Analysis of *var* gene expression in asexual Δ*hdp1* ring-stages found expression of a single major *var* gene, as would be expected for a recently cloned parasite line. This shows that mutually exclusive *var* gene expression remained unaffected in asexual blood stages (fig. S7D) and is consistent with our observation that *hdp1* is not expressed in asexual blood stages and therefore should not alter gene expression in these stages.

**Fig. 3:**
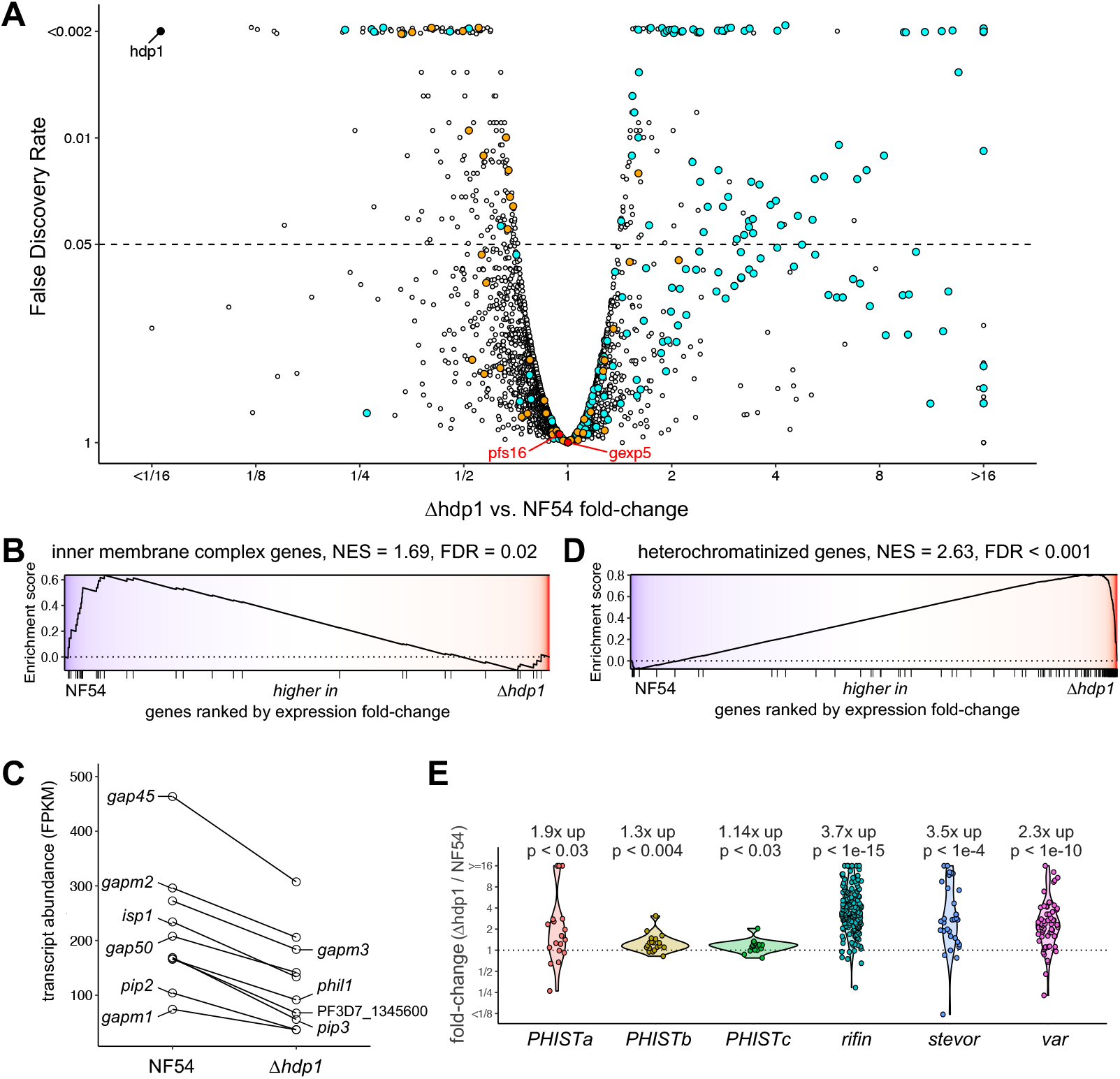
Disruption of HDP1 results in leaky expression of heterochromatin-associated genes and reduced expression of inner membrane complex genes in early gametocytes. **(A)** Genome-wide comparison of differential gene expression in *Δhdp1* and parental NF54 gametocytes on day 2 of gametocytogenesis (stage I, n=2). *hdp1* (solid black), heterochromatin-associated genes (cyan), IMC genes (orange) and the two canonical early gametocyte markers, *pfs16* and *gexp5 (red*) are highlighted. Gene set enrichment analysis (GSEA) indicated significant downregulation of IMC genes **(B-C)** and global upregulation of heterochromatin associated genes **(D-E)**. For the gene enrichment plots (B & D) genes with detectable expression are ranked along the x-axis by expression fold-change, from highest in NF54 to highest in Δhdp1, with tick marks indicating the ranks of inner membrane complex or heterochromatinized genes, respectively. The y-position of the line indicates the enrichment score as a sliding-window for genes that are highest in NF54 to those highest in Δhdp1. The overall normalized enrichment score (NES) and false discovery rate (FDR) for each gene set are shown above each plot. Geometric mean fold-changes and p-values (two-sided, one sample t-test) are indicated in (E).

### HDP1 recognizes a GC-rich DNA motif *in vitro*

Since HDP1 is an integral component of chromatin and has homology to homeo-like domains that typically mediate protein-DNA interactions, we evaluated whether the HDP1 DBD recognizes DNA in a sequence-specific manner using a protein-binding microarray. Recombinant HDP1 DBD was highly enriched on probes containing the palindromic hexamer GTGCAC (Fig. 4A, fig. S8). Since homeo-domains often bind DNA as dimers (*26, 30*), we carried out isothermal titration calorimetry to measure the interaction of HDP1 with double-stranded DNA containing a tandem motif with a 5 bp spacer that places the motifs one helical turn apart. The results found that binding was saturated at a 2:1 protein to DNA molar ratio with a dissociation constant of 2.8 µM, indicating that *in vitro* DNA recognition by the HDP1 DBD occurred primarily as a dimer (Fig. 4B). We subsequently confirmed dimeric binding using DNA gel-shift assays with double-stranded DNA probes containing either no match, a single binding-motif, or a tandem motif. When compared to the tandem-motif probe, the gel-shift of the single motif probe was substantially weaker but identical in size (Fig. 4C), again consistent with DNA recognition as a dimer even when only a single motif is present.

**Fig. 4:**
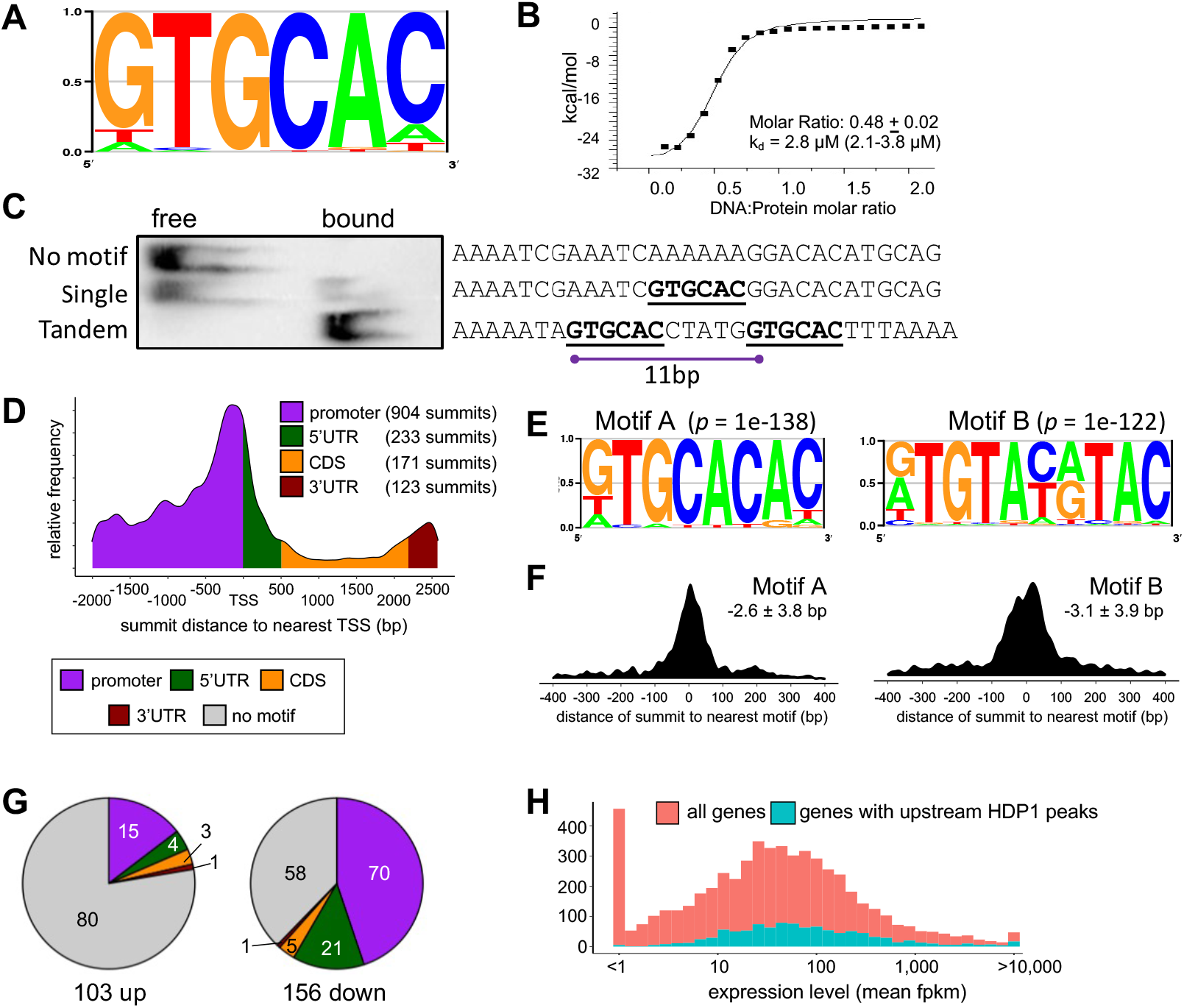
HDP1 binds near the TSS of genes expressed in early gametocytes. **(A)** Maximum enrichment DNA motif for the GST-HDP1 DBD as determined by from protein binding microarray. y-axis shows the fraction of each base at that position. **(B)** Isothermal calorimetry indicates the HDP1-DBD domain recognizes DNA as a dimer. n=2. **(C)** Optimal gel-shift was observed for probes containing a tandem motif with a 5bp spacer compared to probes with either a single or no motif. Representative of n=3. **(D)** Distribution of gene-associated HDP1 ChIP-seq binding sites (n=2). Relative distance to the transcription start site (TSS) is shown on the x-axis. Length of gene features shown is the median length of features with HDP1 binding summits. Summits found within regions mapping to two adjacent genes (242/1188 summits) were counted for both. **(E)** Highly enriched sequence motifs within 100 bp centered on HDP1-GFP ChIP-seq summits. y-axis shows the fraction of each base at that position. **(F)** Density graph of the distance between instances of Motif A (left) or Motif B (right) within HDP1-bound regions to the nearest ChIP-seq summit. The mean distance ± s.e.m. is also shown. **(G)** Number of differentially regulated genes based on location of HDP1-bound regions. **(H)** Histogram of mean expression levels in early gametocytes of NF54 for all genes (red) and genes with upstream (promoter or 5’UTR) HDP1 motifs (teal). Note the lack of genes with HDP1 upstream peaks that have very low expression.

### HDP1 binds GC-rich motifs upstream of a subset of gametocyte-expressed genes *in vivo*

Since our experiments clearly showed that HDP1 can bind DNA and is tightly associated with nuclear chromatin (Fig. 2D), we set out to determine HDP1 genome-wide distribution. To do this we performed chromatin immunoprecipitation sequencing with *anti*-GFP antibodies on nuclei from *hdp1*-GFP and *hdp1-Ty1* gametocytes, with the latter serving as a negative control. We identified 1,003 regions significantly enriched for HDP1-GFP binding containing 1188 binding summits (Data Set 4). Most of these summits (85%) occur within the upstream regions of genes, with the greatest enrichment occurring just upstream of the annotated transcription start site (TSS) (Fig. 4D). 59% of the significantly downregulated genes were found to have upstream HDP1-binding regions, compared to only 18% of those upregulated in Δ*hdp1* gametocytes (Fig. 4G). The prevalence of HDP1-binding near transcription start sites of genes that have reduced expression in gametocytes lacking HDP1 strongly suggest that it functions as a transcriptional activator.

To determine whether HDP1 binding in these regions involved recognition of a specific DNA sequence, we carried out motif enrichment analysis of the 100bp flanking the HDP1 binding summits. This identified two highly enriched sequence motifs, referred to hereafter as Motifs A and B (Fig. 4E). Motif A (GTGCACAC, enrichment *p*-value = 1e-138) is a GC-rich 8mer that closely matched the motif obtained by protein binding microarray (Fig. 4A). Motif B ([GTA]TGTA[CT][GA]TAC, enrichment *p*-value = 1e-122) is a 11mer with greater sequence flexibility that differs substantially from the PBM motif. However, the sequence space covered by the PBM only allows identification of 8bp motifs or shorter. Moreover, ChIP-seq of DNA-binding protein frequently identifies new binding motifs, possibly as the result of interaction with other proteins or domains. Instances of Motif A and B could be identified within 61.8% and 47.8% of HDP1-bound regions, respectively, with 78.1% of HDP1 peaks having at least one motif. The instances within HDP1-bound regions of both Motif A and B occurred were centered on the ChIP-seq summits (Fig. 4F), confirming their recognition by HDP1 *in vivo*. The genome-wide number of these motifs exceeds those found to be occupied by HDP1 indicating that HDP1 is likely recruited to the occupied subset based on interaction with other proteins or excluded from others.

A comparison of expression levels of genes with upstream HDP1-bound regions to all genes, found that HDP1-bound genes are significantly more highly expressed (two-sided Wilcoxon Rank Test *p* = 3.9e-13). HDP1 binding occurred across a wide-range of expression levels but was less frequent upstream of silent or lowly expressed genes (Fig. 4H). Interestingly, while the HDP1 DBD preferentially recognized a tandem motif *in vitro*, virtually all genome-wide binding sites only contain a single motif. Furthermore, the only perfected instances of the tandem motif, which are found at the centromeric end of the subtelomeric repeats showed no significant enrichment, possibly because these regions are not accessible for binding due to heterochromatin formation.

### HDP1 enhances the expression of genes encoding IMC components and required for IMC expansion in early gametocytes

The failure of *Δhdp1* gametocytes to elongate from spherical stage I gametocytes into the oblong Stage II (Fig. 1D) was strikingly similar to the phenotype described for knockdown of PhIL1, an IMC protein that is required for the expansion of the IMC in early *P. falciparum* gametocytes (*31*). Given that genes encoding PhIL1 and other IMC components were highly enriched among genes with reduced expression in HDP1-deficient gametocytes, we examined HDP1 binding at these loci in greater detail. We detected HDP1 binding upstream of eleven out of the 13 IMC genes with reduced expression in Δhdp1 (Fig 5A, fig. S9) and confirmed significant down regulation for all ten genes tested by qRT-PCR (Fig. 5B). Using antibodies against PhIL1 (generous gift of Dr. Pawan Malhotra) we found that PhIL1 expression was greatly reduced in Δ*hdp1* gametocytes (Fig 5C). PhIL1 expression and extension of the IMC during early gametocytogenesis were also clearly impaired upon knockdown of HDP1 (Fig. 5D-E), confirming that both PhIL1 expression in early gametocytes and IMC extension require HDP1.

**Fig. 5:**
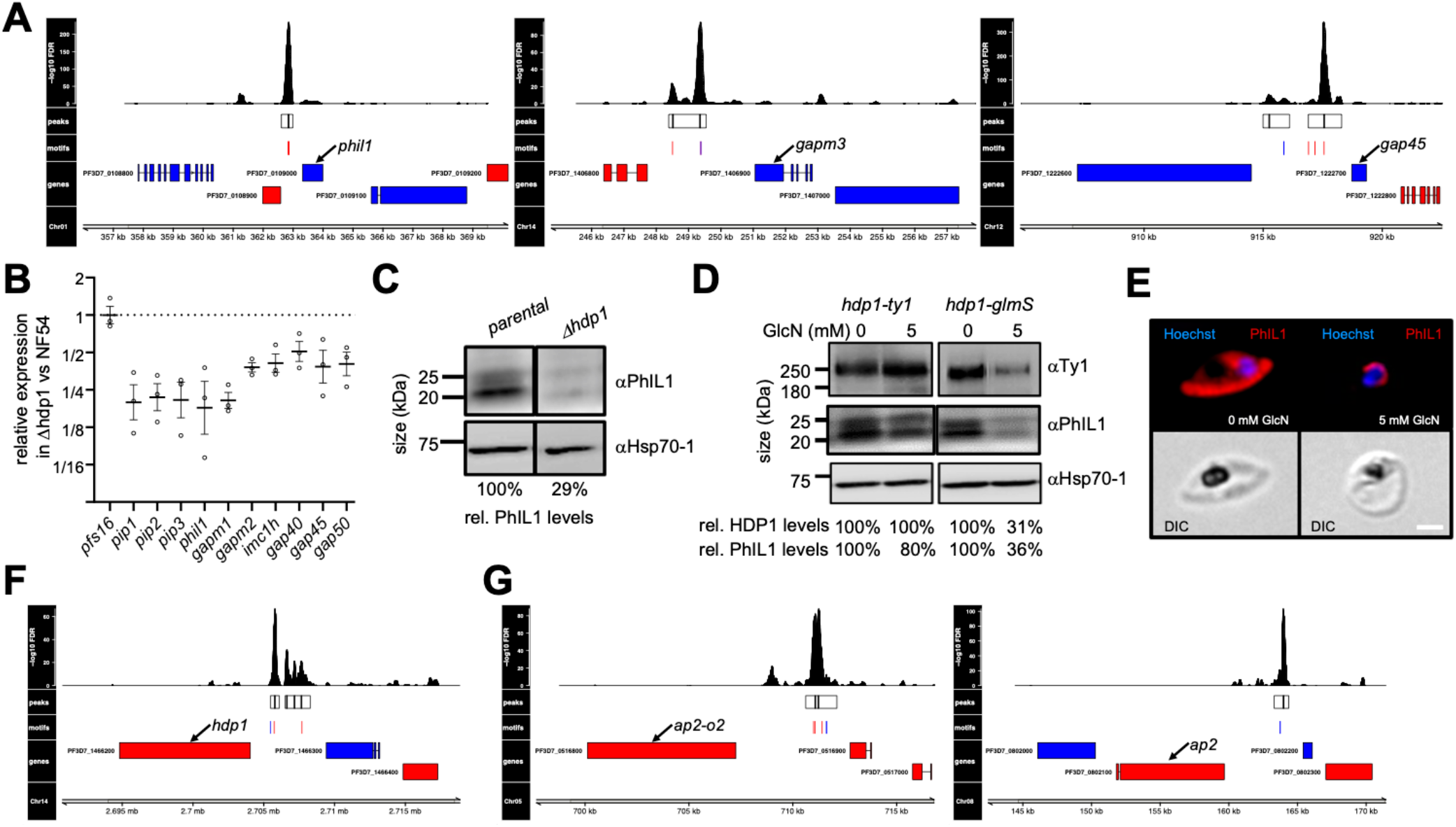
HDP1 is essential for expansion of the inner membrane complex in early gametocytes. **(A)** Example HDP1 binding sites upstream of genes encoding inner membrane complex proteins. Histogram track shows the significance of enrichment by position. Regions of significant enrichment are shown as boxes with black vertical lines indicating peak summits within each peak. Instances of Motif A Motif B, or overlapping motifs within peaks are shown in red, blue and purple, respectively. Genes encoded in forward or reverse orientation are shown in blue or red, respectively. Combined estimate of n=2. **(B)** Validation of down-regulation of genes encoding inner membrane complex genes in HDP1 knockout parasites by qRT-PCR. n=3 **(C)** PhIL1 expression in parental and Δhdp1 gametocytes. Hsp70-1 abundance shown as the loading control. Representative result of n=2. **(D)** Day 5 morphology of *hdp1-ty1* and *hdp1-glmS* gametocytes under 0 and 5 mM glucosamine. HDP1 and PhIL1 protein levels in *hdp1-ty1* and Δ*hdp1* day 5 gametocytes. Hsp70-1 abundance shown as the loading control. Representative result of n=3. **(E)** Immunofluorescence microscopy of PhIL1 distribution in day 5 gametocytes of *hdp1-glmS* under 0 and 5 mM glucosamine. Scale Bar: 3 μm (**F-G)** HDP1 binding sites upstream of genes encoding HDP1 itself and two ApiAP2 proteins. Tracks are the same as in (A).

### HDP1 binds upstream of its own locus and two ApiAp2 genes

Our data indicates that upstream binding of HDP1 promotes the expression of IMC genes necessary for gametocyte maturation. Intriguingly, we also found that the *hdp1* locus itself has multiple upstream HDP1 binding sites (Fig. 5F), pointing to the possibility of a transcriptional feedback loop that may sustain HDP1 expression in gametocytes. This could explain the progressive increase in HDP1 during gametocytogenesis and why the consequences of losing HDP1 expression become apparent when expression is still quite low (Fig. 2C). Additionally, two loci encoding gametocyte-expressed ApiAP2 proteins were found to have HDP1 binding peaks upstream (Fig. 5G) and their expression was significantly reduced in HDP1-deficient gametocytes. These include the gene encoding AP2-O2, which in later stages of gametocytogenesis is required for the upregulation of transcripts essential for ookinete development.

## DISCUSSION

Our understanding of the regulatory machinery that underlies sexual differentiation in *P. falciparum* has improved substantially in recent years. Much of this work has focused on the regulation of AP2-G, the master switch of this developmental decision. However, it is becoming clear that the role of AP2-G is largely constrained to the initiation of the transcriptional program that drives the nearly two week-long process of gametocyte development (*15*). This suggests that a second wave of hitherto unknown transcriptional regulators is required to drive gametocyte-specific gene expression during early gametocyte development.

In this study, we showed that HDP1 is a nuclear DNA-binding protein that functions as regulator of gene expression in early gametocytes and is essential for their development. Our experiments show that HDP1 is an integral component of chromatin in gametocytes and preferentially binds to GC-rich motif near the transcription start site of target genes. Loss of HDP1 expression results in a failure to upregulate a limited set of genes during the early stages of gametocytogenesis, most of which have upstream HDP1 binding sites, supporting HDP1’s role as a positive transcriptional regulator. However, the expression of effected genes was substantially reduced but not completely lost in Δ*hdp1* gametocytes, indicating that upstream binding of HDP1 functions in concert with additional transcriptional regulators to achieve the necessary level of expression.

HDP1 bound upstream of most of the genes being negatively impacted by *hdp1* disruption. Interestingly this Motif A is similar to those recognized by the ApiAP2 proteins SIP2 and API2-I *in vitro* (*36-38*) suggesting that DNA-binding specificity *in vivo* relies on regions beyond the DBD or is mediated through interaction with other proteins (*39*). Motif instances were also found upstream of *hdp1* own locus also indicate the possibility for a positive transcriptional feedback loop that allows HDP1 to enhance its own expression and progressively built-up protein levels. The role of HDP1 in regulating itself and other transcriptional regulators to advance or sustain the transcriptional program underlying gametocytogenesis will certainly warrant additional investigation.

The development of gametocytes lacking HDP1 aborts just prior to the Stage I to Stage II transition, which is marked by the onset of IMC elongation that gives *P. falciparum* gametocytes their eponymous sickle shape. Instead, HDP1-deficient gametocytes remain spherical and lose viability over the next few days. This phenotype closely resembles one described upon knockdown of the IMC protein PhIL1 in early gametocytes. Indeed, we found that HDP1 binds upstream of *phil1* and other genes involved in the expansion of the IMC in early gametocytes and that HDP1 is required for the full expression of these genes. HDP1 also enhances the expression of *mdv1*, another gene known to be essential during early stages of gametocyte maturation (*32*). However, the observed 70% reduction in *mdv1* expression cannot explain the complete arrest of gametocyte maturation upon loss of HDP1, as low-level expression of *mdv1* is sufficient for the normal development of female gametocytes (*32*) while all Δ*hdp1* gametocytes arrest and die during early gametocytogenesis.

While HDP1 is not expressed in *P. falciparum* asexual blood stages and essential during early gametocyte development, whether it is required in other parasite stages remains to be determined. Interestingly, we found that HDP1 enhances the expression of at least two other DNA-binding proteins, indicating that it may also be involved in a cascade of transcriptional regulation that underlies the gene expression changes during late gametocytogenesis and onward. Since the knockdown system used in this study regulates expression at the transcript level, we can infer that *hdp1* mRNA is not required during the later stages of gametocyte maturation but HDP1 protein expressed during the earlier stages of gametocytogenesis may still be required later. Homeodomain-like proteins have been implicated in mating processes of other haploid protozoa, such as Dictyostelium (*34*). As previous transcriptomic studies indicate that HDP1 is also expressed in ookinetes (*35*), HDP1 may well play a role in subsequent stages of the parasite life cycle. Moreover, while expression of HDP1 is gametocyte-specific in the *P. falciparum* blood-stages, disruption of its *P. berghei* ortholog (PBANKA_1329600) significantly impaired asexual blood-stage replication (*33*). This suggests that the gametocyte-specific function of HDP1 may have evolved in the Laveranian clade of malaria parasites, which uniquely produce falciform gametocytes.

Additional studies will be required to elucidate HDP1’s structure-function relationship and identify key interaction partners. Only 2% of its protein sequence is comprised of the homeodomain-like DNA-binding domain while the remainder contains no other identifiable domains and conservation is generally weak. The fact that insertion of a large tag at the N-terminus results in loss of function indicates that critical interactions occur in this region, but which other regions are essential for HDP1’s function remains unclear. Similarly, identifying interactions with other nuclear proteins and genomic locations will offer important insights into its function.

## MATERIALS & METHODS

### Parasite culture

Unless otherwise noted, *P. falciparum* parasites were grown in 0.5% AlbuMAX II supplemented malaria complete medium using stablished cell culture techniques (*40*) at 3% hematocrit and below 3% parasitemia. Strains expressing selectable markers were maintained under constant drug-selection. Toxoplasma tachyzoites were cultured as described in (*41*).

### Gametocyte induction and isolation

Gametocytes were induced synchronously as previously described in (*9*). Gametocyte maturation was monitored by Giemsa-stained thin blood smears and gametocytemia was counted on the fifth day of development. The sexual commitment rate was calculated by dividing the day 5 gametocytemia by the day 1 parasitemia, counted before addition of N-acetyl-D-glucosamine. For knockdown experiments in the HDP1-glmS line, at the gametoring stage either 5 mM glucosamine or solvent control was added. Gametocytes were purified from culture at the required development stage using magnetic columns (LS columns, Miltenyi Biotec).

### Generation of transgenic strains

Transfection of ring-stage parasites were performed as previously described (*42*). Genome editing was performed by CRISR/Cas9 technology using the system described by (*43*). Flanking homology regions were PCR amplified using Advantage Genomic LA polymerase (Takara) from NF54 genomic DNA (for list of primers, see Supplementary Table) and cloned into the *AflII* and *SpeI* sites of pL6 plasmid (carrying hDHFR selectable marker) by Gibson assembly. *Plasmodium* codon optimized sequences for HALO-tag and triple Ty1 epitope tag were synthetized as gene-Blocks (Genewiz). The absence of undesired mutations in the homology regions and the sgRNA was confirmed by Sanger sequencing. Genomic DNA from transfectant parasites was isolated with QIAamp DNA blood Kit (Qiagen, Cat. N° 51106) and diagnostic PCRs were set using Taq Phusion DNA polymerase (Invitrogen). The TGME49_233160-HA parasite line was generated as part of an earlier study by tagging of the endogenous locus in the *T. gondii* RH-ku80ko strain as described in (*41*).

### Flow cytometric analysis of gametocyte viability

Gametocytes were stained with 16 µM Hoechst33342 and 50 nM DilC1 for 30 minutes at 37°C. Using a Cytek DxP12 flow cytometer, gametocytemia was determined by gating for DNA-positive cells and gametocyte viability was inferred based on mitochondrial membrane potential dependent accumulation of DilC1(5) for 1000 gametocytes (*44*). Analysis was carried out using FlowJo 10. The gating strategy is shown in fig. S9.

### Nuclear extract preparation and chromatin high salt fractionation

Nuclear isolation and extraction was carried out based on (*45*), with minor modifications. Briefly, parasites released from RBC’s by saponin treatment (0.01%) were lysed with ice-chilled CLB (20 mM HEPES, pH 7.9; 10 mM KCl; 1mM EDTA, pH 8.0; 1 mM EGTA, pH 8.0; 0.65% NP-40; 1 mM DTT, 1x Roche Complete protease inhibitors cocktail). Nuclei were pelleted at 3,000 x *g* for 20min at 4°C and cytoplasmic fraction was removed. Nuclei were resuspended in digestion buffer (20 mM Tris-HCl, pH 7.5, 15 mM NaCl, 60 mM KCl, 1 mM CaCl_2_, 5 mM MgCl_2_, 300 mM sucrose, 0.4% NP-40, 1 mM DTT, 1x Roche Complete protease inhibitors cocktail EDTA-free) and treated with 5U of micrococcal nuclease (ThermoFisher, Cat. N° 88216) for 30 min in a water bath at 37°C. Soluble and insoluble nuclear fractions were recovered by centrifugation at 3,000 x *g* for 10 min at 4°C. Insoluble nuclear fraction were treated with salt fractionation buffer (10 mM Tris-HCl, pH 7.4; 2 mM MgCl_2_; 2 mM EGTA pH 8.0; 0.1% Triton X-100; 0.1 mM PMSF; 1x Roche Complete protease inhibitors cocktail) supplemented with increasing NaCl concentrations (80-600 mM) while rotating at 4°C for 30 min. All supernatants were recovered by centrifugation at 700 x *g* for 4 min at 4°C and last remaining pellet was resuspended in 1X Phosphate Buffered Saline (PBS) supplemented with protease inhibitors cocktail. 5% of each fraction was prepared for Western blotting to check quality of fractionation.

### Immunoblotting

For SDS-PAGE, total protein lysates were prepared using saponin-lysed parasites resuspended with 1X Laemmli loading buffer diluted in 1x PBS supplemented with 1X Roche Complete protease inhibitors cocktail. Protein samples were separated in 4-15% polyacrylamide gels and transferred to 0.2 µm Immobilion-P^SQ^ transfer membrane (Millipore, Cat. No ISEQ00010) using a Bio-Rad transfer system. Membranes were blocked in 5% skim milk/1x TBS-Tween20 for 1 hour at RT. Primary and secondary antibodies were prepared in 3% skim milk/1x TBS-Tween20 and incubated for 1 hour at RT. Membranes were washed four times with 1x TBS-Tween20 for 10 min, after primary and secondary antibody incubations. The following primary antibodies were used in this study: Anti-Ty1 BB2 mouse (1:2,500; Invitrogen Cat. N^o^ MA5-23513), anti-PhIL1 rabbit (1:5,000 (*46*)), anti-PfHsp70 rabbit (1:5,000; StreesMarq Biosciences Cat. N^o^ SPC-186D), anti-Histone 4 rabbit (1:2,000; Diagenode Cat. N^o^ C15410156-50). HRP-conjugated anti-mouse and anti-rabbit antibodies were used (1:5,000, Millipore). Immunoblots were incubated with the chemiluminescent substrate SuperSignal West Pico PLUS (ThermoFisher, Cat. N^o^ 34578) following manufacturer directions. Chemiluminescent images were obtained using an Azure c300 digital imaging system (Azure Biosystems).

### Live-cell and Immunofluorescence microscopy

For live-cell microscopy of *hdp1-gfp* and NF54 blood stages, infected red blood cells were stained with 16 µM Hoechst33342 in incomplete media for 15 min at 37°C and imaged with identical exposure settings at 1000× magnification using a Leica DMI6000 microscope with differential interference contrast bright field optics, DAPI, and GFP filter cubes. For immunofluorescence microscopy of *hdp1-Ty1* and *hdp1-glmS* gametocytes, cells were immobilized on glass slides with Concanavalin A (5 mg/ml; Sigma) as described in (*47*), then fixed with a solution of 4% paraformaldehyde/0.0075% glutaraldehyde for 20 min at 37°C. Parasites were permeabilized with 0.1% Triton X-100 for 15 min at RT followed by blocking with 3% BSA. Primary antibodies (anti-Ty1 BB2 mouse 1:1,000; anti-PhiL1 rabbit 1:400) were allowed to bind for 1 hour in 3% BSA/PBS followed by three washes with blocking buffer for 5 min. Secondary antibodies were diluted at 1:500 (anti-mouse-Alexa546 and anti-rabbit-Alexa488, Invitrogen) in fresh blocking buffer containing 16 µM Hoechst 33342 and incubated for 1 hour. Z-stacks of stained specimens were collected at 1000× magnification using a Leica DMI6000 microscope with differential interference contrast bright field optics, DAPI, and RFP filter cubes with identical exposure times. Fluorescent channel z-stacks were deconvolved using the ImageJ DeconvolutionLab2 plugin (NLLS algorithm) followed by maximum intensity z-projection and background adjustment. Immunofluorescence microscopy of Toxoplasma tachyzoites was carried out as previously described (*41*).

### Protein Expression and Purification

Expression of recombinant HDP1-DBD motif was done using the Glutathione S-transferase (GST) gene fusion system (GE Healthcare). The pGEX-4T-1 plasmid was used as backbone for cloning the codon optimized sequence comprising the last 87aa of HDP1. Plasmid pGEX-GST-HDP1-DBD was transformed into BL21 (DE3) competent *E. coli* strain (NEB) and protein expression was done following the manufacturer directions with some modifications. Briefly, an overnight culture was inoculated with one bacterial colony in 2X YT media supplemented with the corresponding antibiotic. Next day, culture was diluted 1:100 with fresh media and kept at 30°C with vigorous agitation. Bacterial growth was monitored until culture reach exponential phase. At this point, IPTG (1 mM final concentration) was added, and the culture was kept for another 2 hours at 30°C with vigorous agitation. Cells were harvested and resuspended in lysis buffer (50 mM Tris-HCl, pH 7.5; 100 mM NaCl; 1 mM DTT; 5% Glycerol; 1 mM PMSF; 1 mM EDTA; 1x protease inhibitors cocktail) supplemented with Lysozyme (1mg/ml, Sigma). In order to remove bacterial DNA from our putative DNA binding protein, lysate was treated with polyethyleneimine (PEI) (0.1% v/v) (*48*). Lysate was sonicated, cleared by centrifugation at 14,000 x *g* for 30 min at 4°C. Protein extract was recovered and GST-HDP1-DBD protein was purified using Pierce GST Spin purification kit (Cat. N° 16106) following manufacturer directions. Protein of interest was dialyzed using Slide-A-Lyzer Dialysis Cassette 10,000 MWCO (ThermoScientific, Cat. N° 66810) and concentrated using Amicon Ultra Centrifugal filters 10,000 MWCO (Millipore, Cat. N° UFC901024). Purity was verified by Coomassie staining after SDS-PAGE and concentration was measured by Bradford assay.

### Protein Binding Microarray

GST-HDP1-DBD binding was analyzed twice on Protein Binding Microarrays (PBMs) as previously described (*49, 50*). In this study two different universal PBM arrays (Agilent AMADIDs 016060 v9 and AMADID 015681 v11) were used covering all contiguous 8-mers, as well as gapped 8-mers spanning up to 10 positions. Binding of purified GST-HDP1-DBD fusion proteins was visualized on the PBMs using Alexa-488 conjugated anti-GST antibody. Data analysis was carried out using the PBM analysis software suite downloaded from http://thebrain.bwh.harvard.edu/PBMAnalysisSuite/index.html. Following normalization, enrichment scores were calculated, and the “Seed-and-Wobble” algorithm was applied to the combined data to generate position weight matrices (PWMs). An enrichment score cut-off of 0.45 was used to separate high affinity binding from non-specific and low affinity binding. Secondary motifs were identified by running the “rerank” program until E-scores below 0.45 were obtained.

### Isothermal Titration Calorimetry

Sequence encoding HDP1 aa2991-3078 were cloned into the MCS1 of the pRSFDuet-1 vector (Novagen) engineered with an N-terminal His-SUMO tag. The proteins were expressed in *E. coli* strain BL21 CodonPlus (DE3)-RIL (Stratagene). Bacteria were grown in Luria-Bertani medium at 37°C to OD600=0.8 and induced with 0.4 mM IPTG at 18°C overnight. Cells were collected via centrifugation at 5000×g and lysed via sonication in Lysis Buffer (20 mM Tris-HCl, pH 8.0; 500 mM NaCl; 20 mM imidazole, and 5% Glycerol) supplemented with 1 mM phenylmethylsulfonyl fluoride and 0.5% Triton X-100. Cellular debris was removed by centrifugation at 20,000×g, and the supernatant was loaded onto 5 ml HisTrap FF column (GE Healthcare) and eluted using the lysis buffer supplemented with 500 mM imidazole. The elution was dialyzed at 4°C overnight against the buffer (20 mM Tris-HCl, pH 8.0, 300 mM NaCl, 20 mM imidazole, and 5 mM β-mercaptoethanol) with ULP1 protease added (lab stock). The sample was reloaded on the HisTrap FF column to remove the His-SUMO tag. The flow-through was loaded on the Heparin column (GE Healthcare) and eluted with a gradient of NaCl from 300 mM to 1 M. The target protein was further purified by size exclusion chromatography (Superdex 200 [16/60], GE Healthcare) in the buffer (20 mM Tris-HCl, pH 7.5; 200 mM NaCl; 1 mM MgCl_2_; and 1 mM DTT). The high purity eluting fractions were detected by SDS-PAGE and concentrated to around 20 mg/ml. The protein was flash-frozen in liquid nitrogen and stored at -80°C.

All the binding experiments were performed on a Microcal ITC 200 calorimeter. Purified HDP1-DBD proteins were dialyzed overnight against the buffer (20 mM HEPES, pH 7.5, 150 mM NaCl, 1 mM DTT) at 4°C. DNA oligos were synthesized by Integrated DNA Technologies (IDT) and dissolved in the same buffer. The assays perform with 1 mM DNA duplexes containing the tandem motif (TA<u>GTGCAC</u>CTATG<u>GTGCAC</u>TT) with 0.1 mM HDP1-DBD proteins. Each reaction’s exothermic heat was measured by sequential injection of the 2 µL DNA duplexes into proteins solution, spaced at intervals of 180 seconds. The titration was according to standard protocol at 20°C and the data were fitted using the program Origin 7.0.

### Gel-shift Assays

Electrophoretic mobility shift assays were performed using Light Shift EMSA kits (Thermo Scientific) using 24 pg of protein and 40 fmol of probe, as previously described (*37*). Biotinylated double-stranded were synthesized probes (ThermoFisher) with the indicated sequence.

### RNA Extraction, cDNA synthesis, and quantitative RT-PCR

Total RNA from saponin-lysed parasites was extracted using Trizol (Invitgrogen) and Direct-Zol RNA MiniPrep Plus kit (Zymo Research). The cDNA was prepared from 100-500ng total RNA (pre-treated with 2U DNase I, amplification grade) using SuperScript III Reverse Transcriptase kit (Invitrogen) and random hexamers. Quantitative PCR was performed on the Quant Studio 6 Flex (Thermo Fisher) using iTaq Sybr Green (Bio-Rad) with specific primers for selected target genes (Data set S5). Quantities were normalized to seryl-tRNA synthetase (PF3D7_0717700). Analysis of expression of the *var* gene family was performed by using the primer set described in Salanti et al. 2003 (*51*).

### RNA sequencing

Following gametocyte induction, highly synchronous cultures containing committed schizonts were added to fresh RBCs and allowed to reinvade for 12 hours prior to the addition of 50 mM N-acetyl glucosamine to block the development of hemozoin-containing asexual trophozoites. On day 2 of gametocyte development, stage I gametocytes were purified magnetically, and total RNA was extracted as described above. Following RNA isolation, total RNA integrity was checked using a 2100 Bioanalyzer (Agilent Technologies, Santa Clara, CA). RNA concentrations were measured using the NanoDrop system (Thermo Fisher Scientific, Inc., Waltham, MA). Preparation of RNA sample library and RNA-seq were performed by the Genomics Core Laboratory at Weill Cornell Medicine. rRNA was removed from Total RNA using Illumina Ribo Zero Gold for human/mouse/rat kit. Using Illumina TruSeq RNA Sample Library Preparation v2 kit (Illumina, San Diego, CA), Messenger RNA was fragmented into small pieces using divalent cations under elevated temperature. The cleaved RNA fragments were copied into first strand cDNA using reverse transcriptase and random primers. Second strand cDNA synthesis followed, using DNA Polymerase I and RNase H. The cDNA fragments then went through an end repair process, the addition of a single ‘A’ base, and then ligation of the adapters. The products were then purified and enriched with PCR to create the final cDNA library. Libraries were pooled and sequenced on Illumina HiSeq4000 sequencer with single-end 50 cycles. Read files were checked for quality by using FASTQC v0.11.5 (https://github.com/s-andrews/FastQC). Reads were trimmed to remove low-quality positions and adapter sequences using cutadapt (version 1.16) (*52*). Reads were mapped against the *P. falciparum* 3D7 reference genome v40 (*53*) using STAR aligner (version 2.61) (*54*) and nuclear-encoded genes were analyzed for differential gene expression using cufflinks (version 2.2.1) (*55*). Genes with a false discovery rate of <=0.05 with a mean FPKM >5 in at least one strain were called significant. For genes with FPKM > 5 in one strain and no detectable expression in the other, FKPM values were set to 0.1 for purposes of fold-change calculation. Gene Set Enrichment Analysis was carried out with the FGSEA v1.16.0 Bioconductor v 3.12 package (*56*) with an FDR cutoff of <=0.05.

### Chromatin-Immunoprecipitation Sequencing (ChIP-seq)

*hdp1-gfp* and *hdp1-Ty1* gametocytes were harvested on day 5 of development, isolated from the middle interface of a 30/35/52.5% percoll gradient, washed in 1X PBS, and then spun down for 5 min at 2,500 x *g*. Pelleted parasites were resuspended in 500 μL of lysis buffer (25 mM Tris-HCl, pH 8.0, 10 mM NaCl, 2 mM AESBF, 1% NP-40, 1X protease inhibitors cocktail) and incubated for 10 min at RT. Parasite lysates were homogenized by passing through a 26G ½ needle 15 times. Samples were crosslinked by adding formaldehyde to a final percentage of 1.25% followed by further homogenization by passing through a 26G ½ needle 10 times and incubation for 25 min at RT while continuously mixing. Crosslinking was quenched by adding glycine to a final concentration of 150 mM followed by incubation for 15 min at RT and then 15 min at 4°C with continuous mixing. Samples were spun for 5 min at 2,500 x *g* at 4°C and crosslinked parasite pellets were washed once with 500 μL ice-cold wash buffer (50 mM Tris-HCl, pH 8.0, 50 mM NaCl, 1 mM EDTA, 2 mM AESBF, 1x protease inhibitors cocktail). Samples were stored at -80°C until further use. The crosslinked parasite pellets were thawed on ice and resuspended in 1 mL of nuclear extraction buffer (10 mM HEPES, 10 mM KCl, 0.1 mM EDTA, 0.1 mM EGTA, 1 mM DTT, 0.5 mM AEBSF, 1X protease inhibitor cocktail). After a 30 min incubation on ice, Igepal-CA-630 was added to a final concentration of 0.25% and homogenized by passing through a 26G x ½ needle 7 times. The nuclear pellet was extracted by centrifugation at 2,5000 x *g*, then resuspended in 130 µl of shearing buffer (0.1% SDS, 1 mM EDTA, 10 mM Tris-HCl pH 7.5, 1X protease inhibitor cocktail) and transferred to a 130 µl Covaris sonication microtube. The sample was then sonicated using a Covaris S220 Ultrasonicator for 8 min (Duty factor: 5%, Intensity peak power: 140, Cycles per burst: 200, Bath temperature: 6°C). Samples were transferred to ChIP dilution buffer (30 mM Tris-HCl pH 8, 3 mM EDTA, 0.1% SDS, 300 mM NaCl, 1.8% Triton X-100, 1X protease inhibitor cocktail, 1X phosphatase inhibitor tablet) and centrifuged for 10 min at 16,000 x *g* at 4°C, retaining the supernatant. For each sample, 13 μl of protein A agarose/salmon sperm DNA beads were washed three times with 500 µl ChIP dilution buffer (without inhibitors) by centrifuging for 1 min at 800 x *g* at room temperature, then buffer was removed. For pre-clearing, the diluted chromatin samples were added to the beads and incubated for 1 hour at 4°C with rotation, then pelleted by centrifugation for 1 min 800 x *g*. Supernatant was removed carefully so as not to remove any beads. 10% of the sample was removed to be used as input, and 2 µg of anti-GFP antibody (Abcam ab290, anti-rabbit) were added to the remaining sample and incubated overnight at 4°C with rotation. For each sample, 25 µl of protein A agarose/salmon sperm DNA beads were washed with ChIP dilution buffer (no inhibitors), blocked with 1 mg/mL BSA for 1 hour at 4°C, then washed three more times with buffer. 25 µl of washed and blocked beads were then added to the sample and incubated for 1 hour at 4°C with continuous mixing to collect the antibody/protein complex. Beads were pelleted by centrifugation for 1 min at 800 x *g* at 4°C. The bead/antibody/protein complex was then washed for 15 min with rotation using 1 mL of each of the following buffers twice: low salt immune complex wash buffer (1% SDS, 1% Triton X-100, 2 mM EDTA, 20 mM Tris-HCl pH 8, 150 mM NaCl), high salt immune complex wash buffer (1% SDS, 1% Triton X-100, 2 mM EDTA, 20 mM Tris-HCl pH 8, 500 mM NaCl), LiCl immune complex wash buffer (0.25M LiCl, 1% Igepal, 1% sodium deoxycholate, 1 mM EDTA, 10 mM Tris-HCl pH 8), and TE wash buffer (10 mM Tris-HCl pH 8, 1 mM EDTA). The complex was then eluted from the beads by adding 250 μl of freshly prepared elution buffer (1% SDS, 0.1 M sodium bicarbonate), twice with 15 min rotation. We added 5 M NaCl to the elution and cross-linking was reversed by heating at 45°C overnight followed by addition of 15 μl of 20 mg/mL RNase A with 30 min incubation at 37°C. After this, 10 μl 0.5 M EDTA, 20 μl 1 M Tris-HCl pH 7.5, and 2 μl 20 mg/mL proteinase K were added to the elution and incubated for 2 hours at 45°C. DNA was recovered by phenol/chloroform extraction and ethanol precipitation, using 1 volume of phenol/chloroform/isoamyl alcohol (25:24:1) mixture twice and chloroform once, then adding 1/10 volume of 3 M sodium acetate pH 5.2, 2 volumes of 100% ethanol, and 1/1000 volume of 20 mg/mL glycogen. Precipitation was allowed to occur overnight at -20°C. Samples were centrifuged at 16,000 x *g* for 30 min at 4°C, then washed with fresh 80% ethanol, and centrifuged again for 15 min with the same settings. Pellet was air-dried and resuspended in 50 μl nuclease-free water. DNA was purified using Agencourt AMPure XP beads. Libraries were then prepared from this DNA using a KAPA library preparation kit (KK8230) and sequenced on a NovaSeq 6000 machine.

### Analysis of ChIP-seq Data

Reads in sequenced libraries were checked for quality with FASTQC v0.11.9 (https://github.com/s-andrews/FastQC), adapter- and quality-trimmed with trimmomatic v0.39 (*52*). Properly paired reads were then aligned against the *P. falciparum* 3D7 genome v51 with bwa v0.7.17 (*57*) and sorted by name using samtools v1.13 (*58*). Differential genome-wide enrichment between the HDP1-GFP and HDP1-Ty1 samples was then calculated using macs2 v2.2.7.1 (*59*) to generate poisson-based FDR and fold-enrichment tracks and significant peaks. Peak annotation and visualization were carried out in R using the GenomicRanges v1.44.0 (*60*), Gviz v1.36.2 (*61*), and ChIPpeakAnno v3.26.2 (*62*) packages from Bioconductor (*63*) using *P. falciparum* 3D7 genome annotation v51 from PlasmoDB. Gene promoters were defined as 2kb upstream of the annotated TSS and “upstream region” is defined as the promoter + 5’UTR. Motif enrichment for 100bp centered on HDP1-GFP ChIP-seq summits was carried out using the HOMER2 v4.11 findMotifsGenome function (*64*).

## Supporting information

Supplementary Data Sets

## General

We wish to thank Dr. Pawan Malhotra for the generous gift of anti-PhIL1 antibodies,

V. Carruthers for the generous gift of anti-Ty1 antibodies, the Weill Cornell Medicine genomics core for technical support, respectively, as well as K. Deitsch and J. King for valuable feedback on the manuscript.

## Author contributions

Conceptualization: B.F.C.K.; Methodology: B.F.C.K., R.C.M., W.X., K.G.L.; Investigation: R.C.M., X.T., W.X., L.O., W.D.; Software, Formal Analysis, Data Curation: B.F.C.K.; Writing – Original Draft: R.C.M.; Writing – Review & Editing: B.F.C.K., R.C.M.; Visualization: R.C.M., B.F.C.K.; Supervision: B.F.C.K., K.G.L., M.L., D.P.; Project Administration: B.F.C.K; Funding Acquisition: B.F.C.K.

## Data and materials availability

Raw high throughput sequencing data have been deposited in the NCBI Sequence Read Archive under accession number SRPXXXX.

Analysis code is available at https://github.com/KafsackLab/HDP1.

All data needed to evaluate the conclusions in the paper are present in the paper and/or the Supplementary Materials. Additional data related to this paper may be requested from the authors.

**fig. S1:**
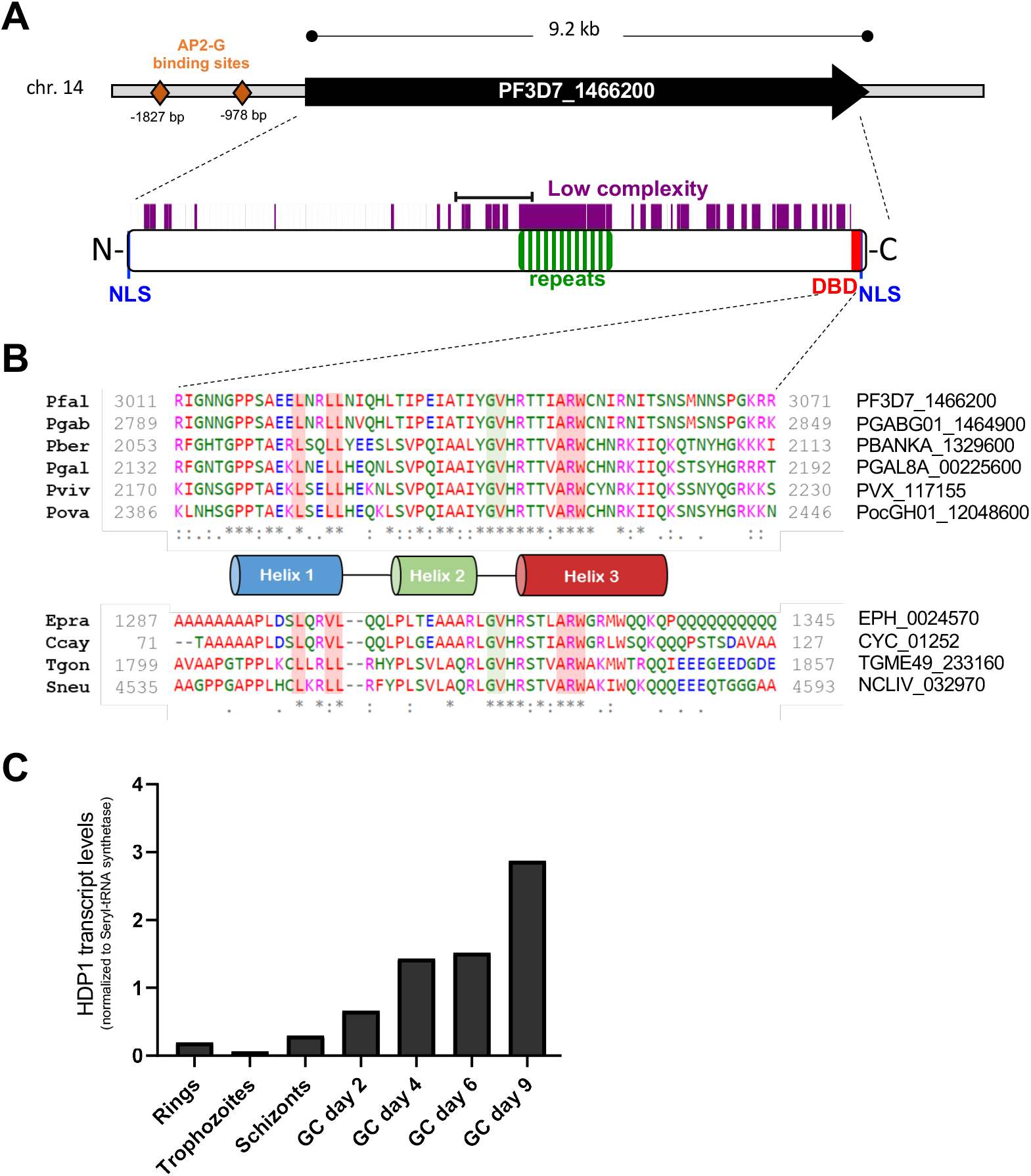
The predicted DNA-binding protein HDP1 is expressed in gametocytes. **(A)** The single exon locus Pf3D7_1466200 encodes a large 3078aa protein with a predicted C-terminal Helix-Turn-Helix DNA-binding domain (DBD) and two nuclear localization signals (NLS). Multiple AP2-G binding sites are found in its 2kb promoter region. Black bracket indicates the antigen used for generating antisera used in (29). **(B)** Alignment of the helix-turn-helix domain for homologs from other apicomplexan parasites. **(C)** Quantitative RT-PCR of *hdp1* transcripts found minimal expression in asexual blood stages with upregulation during gametocyte development. (mean of n=2). Transcript levels were normalized to normalized to Ser-tRNA synthase.

**fig. S2:**
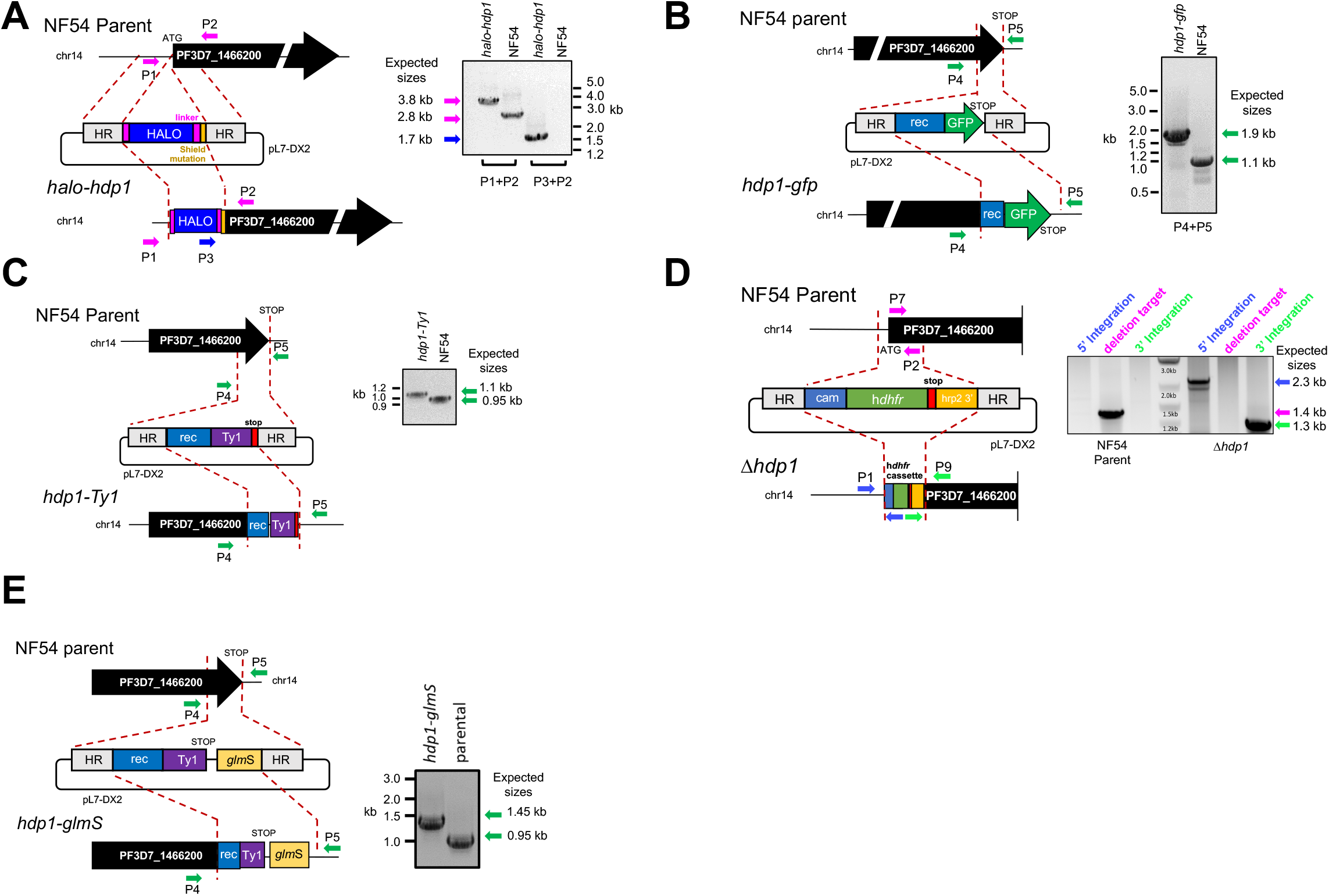
Validation of engineered parasite lines. **(A)** Generation of h*alo-hdp1* parasites by CAS9 genome editing. Insertion of the N-terminal HALO tag at the 5 ’end the *hdp1* coding sequence was confirmed by PCR and checked for mutations by Sanger sequencing of the 3.8kb PCR product (not shown). **(B)** Generation of *hdp1-gfp* parasites by CAS9 genome editing. Insertion of the C-terminal GFP tag at the 3 ’end the *hdp1* coding sequence was confirmed by PCR and checked for mutations by Sanger sequencing of the 1.9kb PCR product (not shown). **(C)** Generation of *hdp1-Ty1* parasites by CAS9 genome editing. Insertion of the C-terminal triple Ty1 epitope tag at the 3’ end the *hdp1* coding sequence was confirmed by PCR and checked for mutations by Sanger sequencing of the 1.1kb PCR product (not shown). **(D)** Generation of *Δhdp1* parasites by CAS9 genome editing. Replacement of 1.4 kb flanking the *hdp1* start codon by a hDHFR selectable marker cassette was confirmed by PCR. **(E)** Generation of *hdp1-glmS* parasites by CAS9 genome editing. Insertion of the C-terminal triple Ty1 epitope tag and the *glmS* ribozyme at the 3’ end the *hdp1* coding sequence was confirmed by PCR and checked for mutations by Sanger sequencing of the 1.4 kb PCR product (not shown).

**fig. S3:**
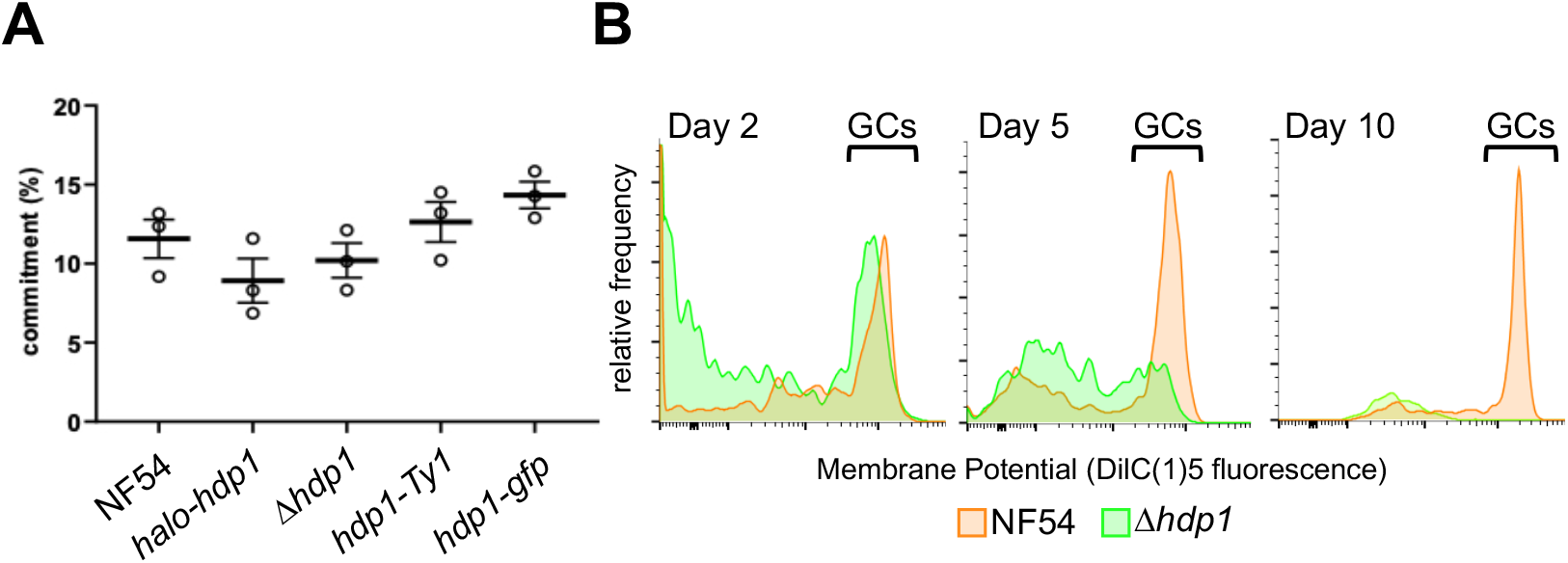
Loss of HDP1 does not alter the sexual commitment frequency or Stage I gametocyte viability. **(A)** The sexual commitment frequency (day 5 gametocytes per day 1 ring stages) is not significantly affected in *halo-hdp1* and *Δhdp1* parasites. n=3 **(B)** Mitochondrial membrane potential of gametocytes (GCs), as measured by DilC(1)5 staining, indicates similar viability on day 2, but not days 5 or 10, for NF54 (orange) and Δ*hdp1* (green) gametocytes. Representative of n=2.

**fig. S4:**
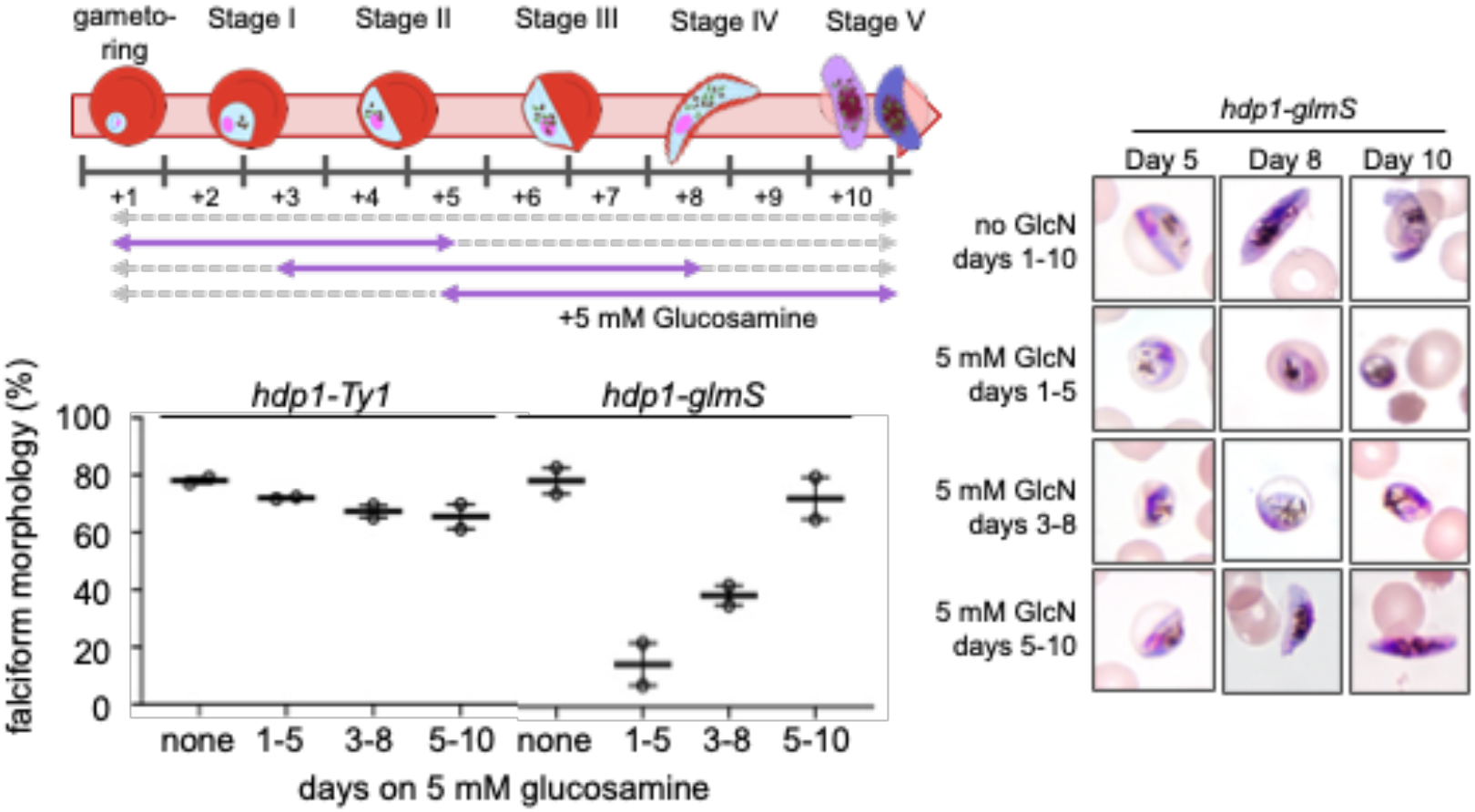
Inducible knockdown of HDP1 reduces gametocyte maturation in early but not late gametocytes. Representative morphology (right) of *hdp1-glmS* gametocytes in response to 5 mM glucosamine on days 1-5, 3-8, 5-10, or in the absence of glucosamine. Percentage of falciform gametocytes on Day 10 in response to 5 mM glucosamine on days 1-5, 3-8, 5-10, or in the absence of glucosamine for *hdp1-Ty1* or *hdp1-glmS* parasites (bottom). mean±s.e.m of n=2.

**fig. S5:**
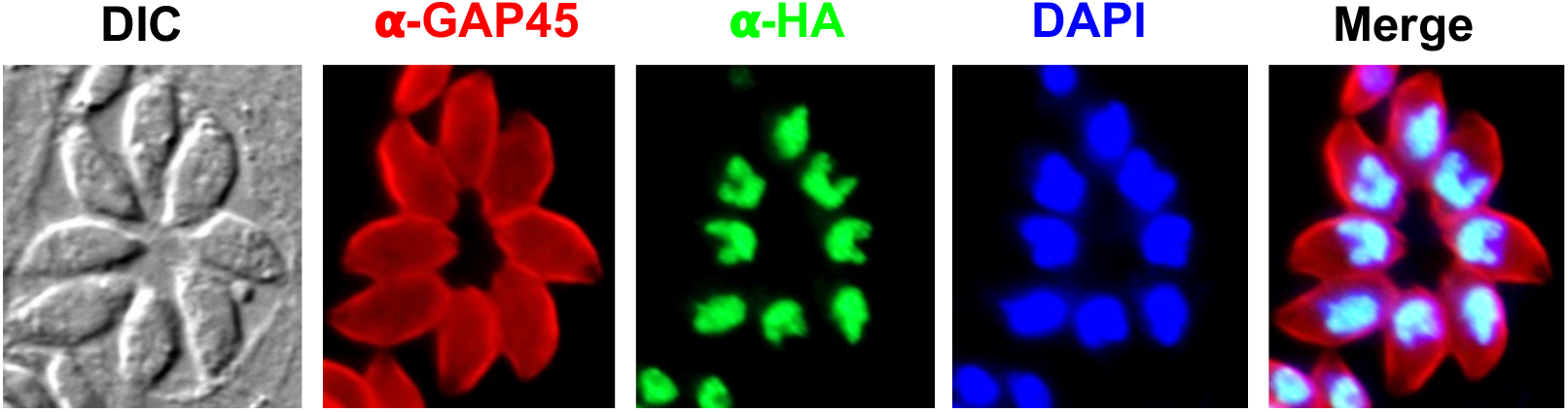
Immunofluorescence microscopy localizes the HA-tagged ortholog TGME49_233160 to the nucleus of *Toxoplasma gondii* tachyzoites.

**fig. S6:**
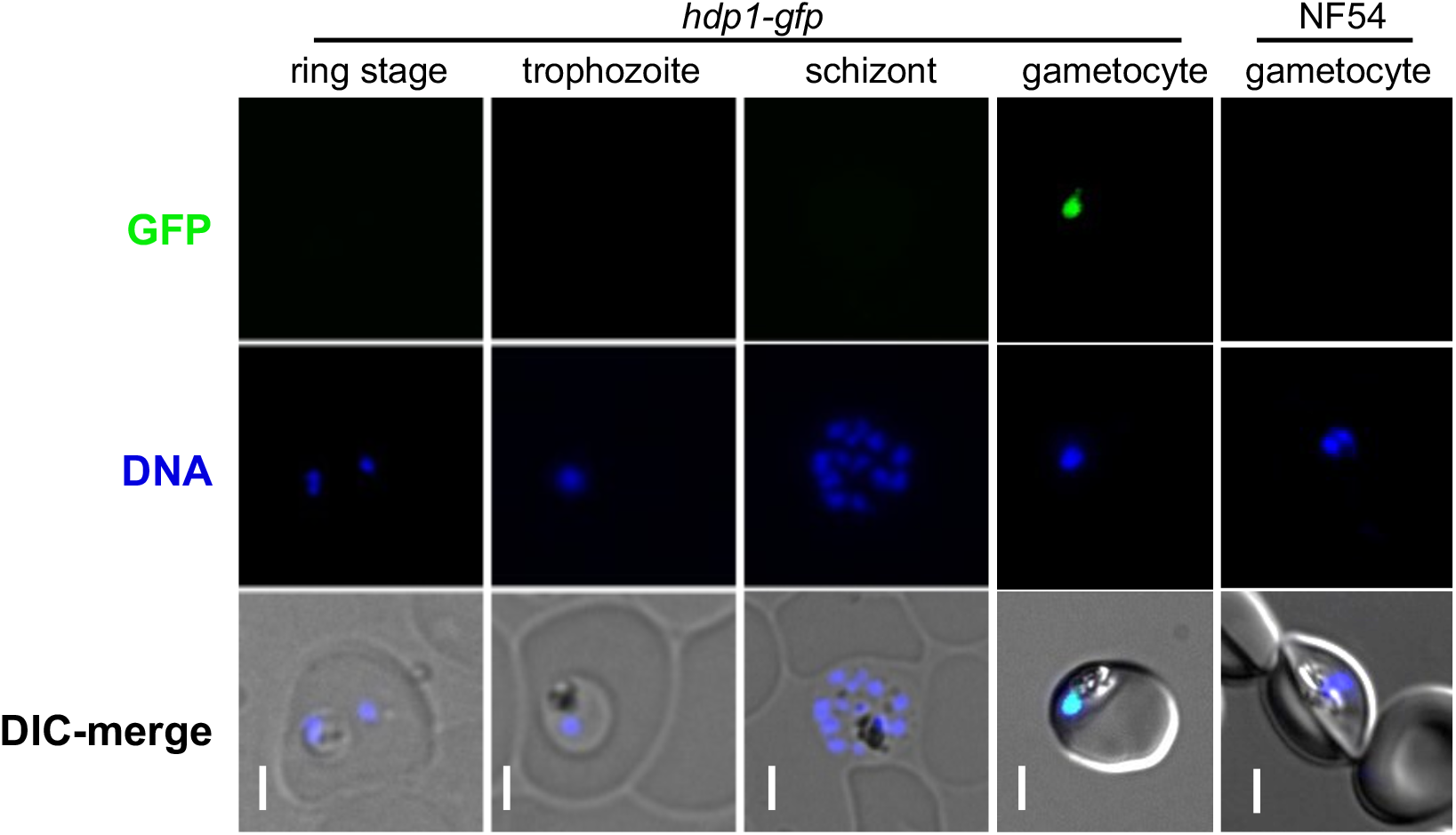
HDP1-GFP localized to the nucleus of *hdp1-gfp* gametocytes. No signal was observed in *hdp1-gfp* asexual blood stages or gametocytes of the untagged NF54 parent line. Scale bar is 3 microns. Exposure and brightness/contrast settings are uniform across the images shown. Representative of n=3.

**fig. S7:**
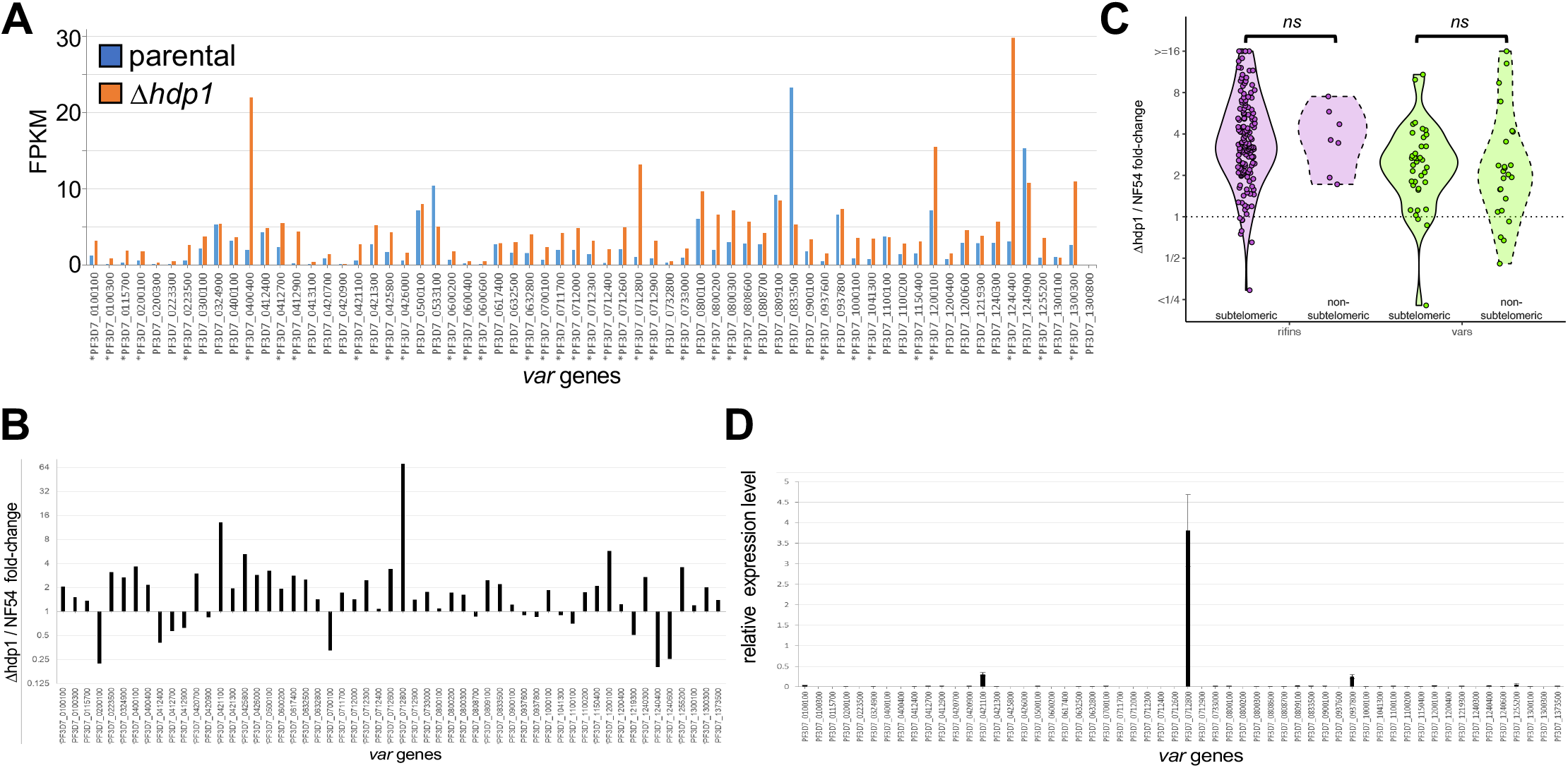
*var* gene expression is altered in early Δ*hdp1* gametocytes but not asexual blood stages. **(A)** Normalized abundance of reads uniquely mapping to *var* genes in day 2 gametocytes from Δ*hdp1* (orange) or parental NF54 parasites (blue). Significantly upregulated genes are marked with asterisks. (n=2) **(B)** qRT-PCR confirmation of *var* gene up-regulation in Δ*hdp1* vs NF54 day 2 gametocytes. **(C)** Upregulation of *rifin* (purple) and *var* genes (green) in *Δhdp1* day 2 gametocytes is independent of chromosomal position in subtelomeric or non-subtelomeric heterochromatin clusters. **(D)** qRT-PCR analysis of *var* transcript abundance (normalized seryl tRNA synthetase expression) in Δ*hdp1* asexual ring stages found expression a single dominant var as expected indicating that mutually exclusive expression remains unchanged in asexual stages. (n=2)

**fig. S8:**
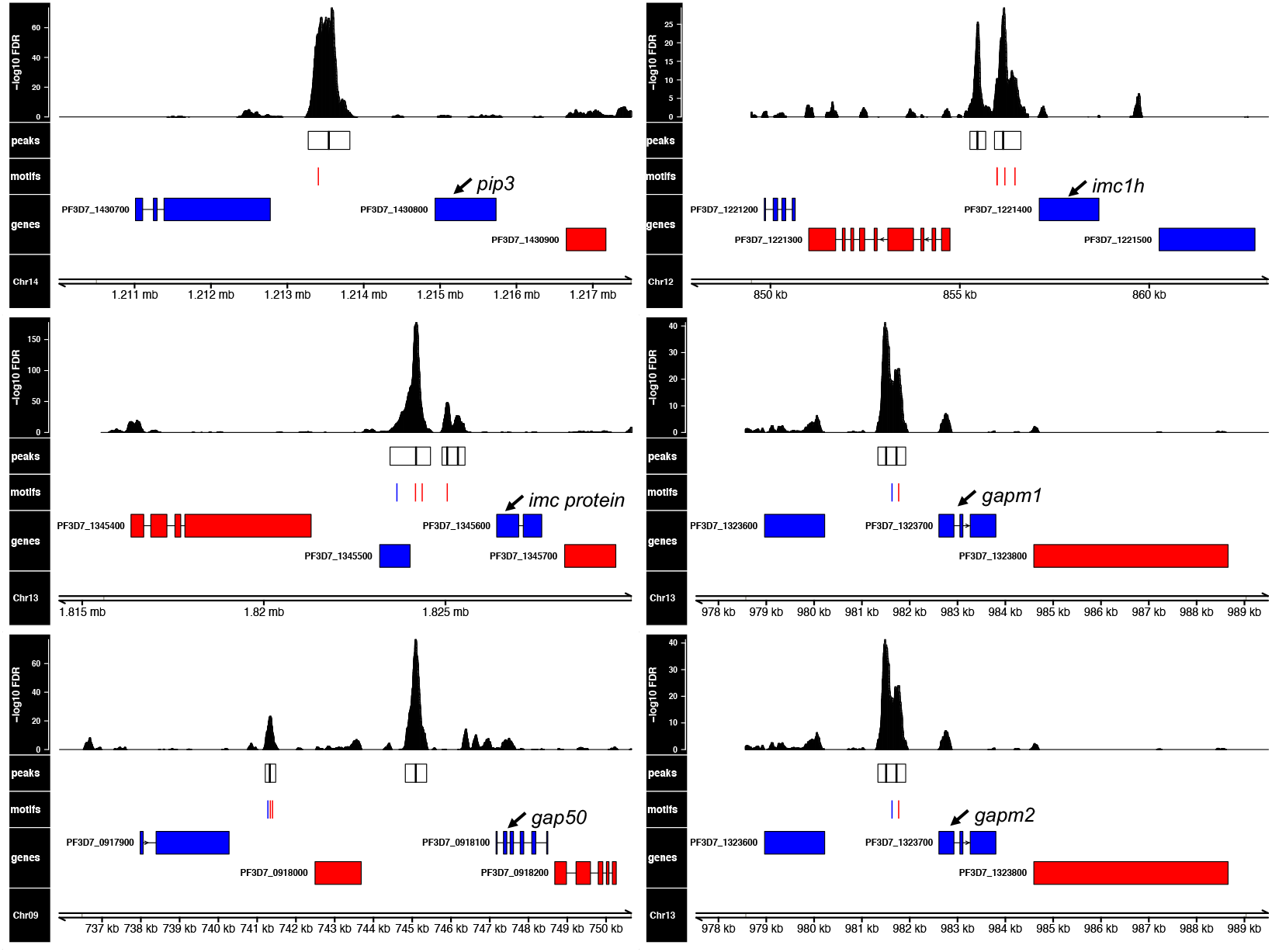
Genes encoding inner membrane complex genes with significantly reduced expression in HDP1 knockout parasites have upstream HDP1 binding sites. Remaining HDP1 binding sites upstream of genes encoding inner membrane complex proteins. Histogram track shows the significance of enrichment by position. Regions of significant enrichment are shown as boxes with black vertical lines indicating peak summits within each peak. Instances of Motif A, Motif B, or overlapping motifs within peaks are shown in red, blue and purple, respectively. Genes encoded in forward or reverse orientation are shown in blue or red, respectively. Combined estimate of n=2.

**fig. S9:**
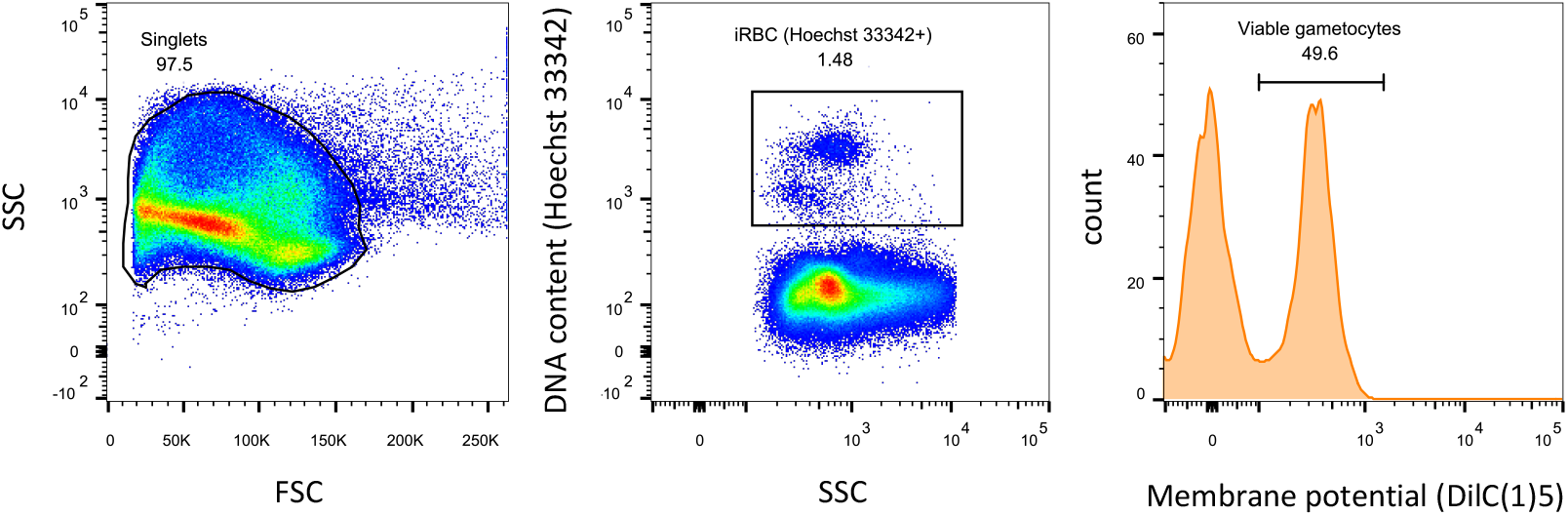
Gating schema for viable gametocytes. Populations were gated for single cells based on forward (FSC) and side scatter (SSC). Viable gametocytes were identified based on DNA content and mitochondrial membrane potential based on Hoechst33342 and DilC(1)5 staining.

